# Distinct control of T cell proliferation and effector function by partitioning of intracellular sulfur from cysteine

**DOI:** 10.64898/2026.01.27.702014

**Authors:** Beth Kelly, Minsun Cha, Tatjana Gremelspacher, Jacob L. Martin, Massimo Andreis, Gustavo E. Carrizo, Mia Gidley, Michal A Stanczak, Petya Apostolova, David E. Sanin, Ananya Majumdar, Erika L. Pearce

## Abstract

Delineating how acquired nutrients are partitioned into different intracellular pathways, and how these various fates support distinct functions in T cells is limited. We show that CD8^+^ T cells acquire cysteine to serve both as a substrate for glutathione (GSH) production, which modulates effector functions, and to cede its sulfur for NFS1-dependent FeS-cluster synthesis, which supports proliferation. NFS1 deletion in activated CD8^+^ T cells promotes exhaustion and dampens anti-cancer immunity, while blocking cysteine flux into GSH, or enforcing FeS metabolism, enhance tumor control. This role for disrupted FeS metabolism in T cell exhaustion is echoed in data from human HCC. Elucidating how different intracellular pathways use cysteine enables targeted control of cysteine flux to retain beneficial effects of cysteine while abolishing those that restrain function. We illustrate this concept for one metabolite, cysteine, but it is likely to apply to other metabolites relevant for immune cell function.

## INTRODUCTION

CD8^+^ T cell activation in response to cognate antigen and costimulation is accompanied by cell growth, extensive proliferation, and acquisition of effector functions, including cytolytic capacity and expression of cytokines like interferon (IFN)γ and tumor necrosis factor (TNF). CD8^+^ T cells are critical for fighting cancer and infection, yet they sometimes fail to control disease. Inadequate T cell responses manifest in many ways, e.g. weak antigenicity, poor tissue infiltration, or T cell exhaustion^1–3^. Why certain cancers evade immune control is not completely understood, but often can in part be attributed to insufficient T cell function. To acquire appropriate function, T cells tune their metabolism to support the demands of activation, differentiation, proliferation, and effector molecule expression^4^.

Like carbohydrates and lipids, amino acids are fundamental building blocks supporting life. While their use for protein synthesis is well-established, individual amino acids have many other critical roles, including in energy metabolism, signaling, and maintenance of redox balance^5^. The 9 essential amino acids (EAA) must be acquired from dietary sources, while the remaining nonessential amino acids (NEAA) can be synthesized to sufficient levels in the body. However, NEAA can become essential when cellular demand outstrips synthesis, and cells must then acquire these from extracellular sources^5,6^. Many physiological contexts in which NEAA become essential remain to be defined.

Many immunometabolism studies have focused on the supply or early catabolism of a particular nutrient, and less on the fact that a single nutrient can contribute to multiple intracellular pathways, each potentially having a different effect on function. In our previous work studying a eukaryotic slime mold that undergoes unicellular proliferation versus multicellular development depending on food availability, we showed that reactive oxygen species (ROS), generated as a consequence of nutrient limitation, led to the sequestration of cysteine in the antioxidant GSH^7^. The pull of available cysteine into GSH limited the use of cysteine’s sulfur atom in FeS cluster synthesis, which supports mitochondrial respiration and cell proliferation, as many enzymes, including several in the ETC, depend on FeS complexes for activity^8^. This regulated sequestration of sulfur upon nutrient fluctuation dictated cell, and thus organismal, fate, by forcing a switch from unicellular proliferation to nonproliferating multicellular aggregation. Sulfur metabolic machinery is highly conserved in eukaryotes^9^, indicating evolutionary pressure to preserve sulfur metabolism, and suggesting that differential intracellular routing of sulfur, influenced by its availability, will have profound consequences in mammalian cells. With this in mind, we set out to establish the molecular mechanisms of sulfur utilization, which is supplied to our cells by methionine and cysteine, and how sulfur itself contributes to T cell function.

## RESULTS

### Cysteine starvation of activated T cells enhances cytokine production but impairs proliferation

Rather than assess sulfur depletion during T cell activation, we sought to determine its effect on fully activated effector CD8^+^ T cells, reasoning that later sulfur limitation would more closely resemble the fate of activated T cells as they leave the nutrient supportive lymph node and traffic to tissue sites. CD8^+^ T cells isolated from naïve mice were activated in vitro with αCD3/28 + IL-2 for 48 h, cultured for a further 3 days in IL-2, and then placed in medium without methionine, cysteine, without both methionine and cysteine (sulfur amino acids, SAA), or without glutamine for 6 – 24 h (Figure 1A). Methionine is an EAA and must be acquired from the diet, while cysteine is considered a NEAA as it can be made from methionine, but can become conditionally essential depending on cellular demand^5,6^. Cell viability decreased after 24 h without methionine or SAAs (Figure S1A). 24 h cysteine starvation resulted in intracellular cysteine depletion (Figure 1B), with no viability loss (Figure 1C); cells did not lose viability until 3 days of cysteine starvation (Figure S1B). As cells were viable after 6 h starvation in all conditions, we measured interferon (IFN)γ production after PMA/ionomycin restimulation. 6 h cysteine-starved cells had comparable IFNγ expression to controls, while cells starved of methionine, SAAs, or glutamine (a non-sulfur-containing AA) had decreased IFNγ (Figure S1C, S1D). However, after 24 h (when cysteine-starved cells were still viable, but methionine-starved cells were not), cysteine-starved T cells expressed more IFNγ and tumor necrosis factor (TNF) upon restimulation (Figure 1D, 1E). OTI-transgenic CD8^+^ T cells (ovalbumin (OVA)-specific) depleted for cysteine had enhanced cytotoxicity, causing increased death of EL4 lymphoma cells expressing OVA (EL4-OVA) after 6 hours of co-culture (Figure 1F, G). 6 h starvation of EL4-OVA cells did not impair their viability, in the absence of OT-I T cell co-culture (Figure S1E), indicating that the increased EL4-OVA cell death observed when co-cultured with cysteine-starved OT-I T cells was due to improved killing activity of the T cells. Despite this enhanced functionality, cysteine-starved T cells had dampened proliferation, indicated by a lower percentage of cells positive for EdU and FxCycle violet staining (Figure 1H). Specifically, cysteine-starved T cells were stalled in the G0/G1 phase of cell division, with fewer cells in the S and G2/M phases compared to fully fed cells (Figure 1I). Proliferation was also inhibited in other starvation conditions after 6 h, prior to lost viability (Figure S1F).

**Figure 1.**
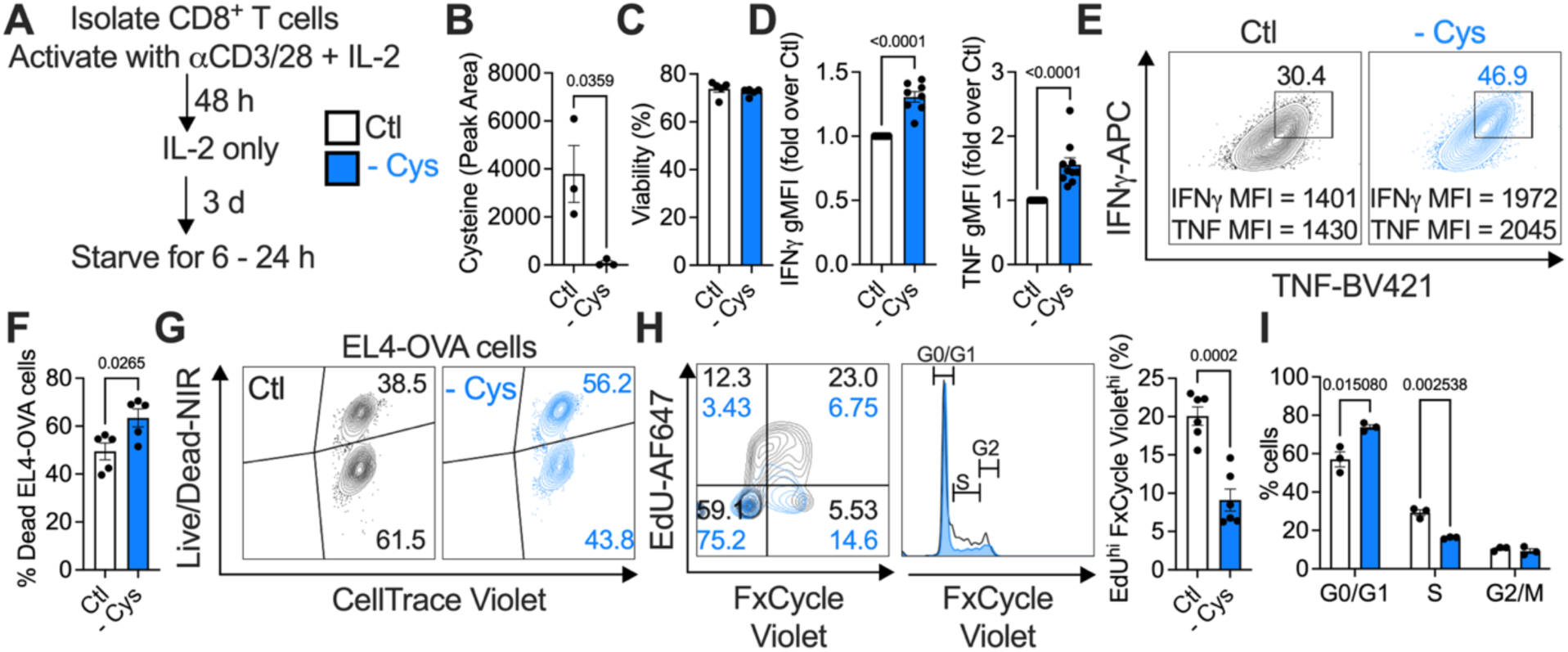
Cysteine starvation enhances CD8^+^ T cell cytokine production but impairs proliferation. (A - I) Isolated, activated (with αCD3/28) CD8^+^ T cells were starved of cysteine for 6 - 24 h (from day 5 to day 6 of culture). (B) Cysteine depletion in starved cells, measured by LC-MS. (C) viability, and (D, E), IFNγ and TNF production in fed and starved cells, restimulated for 5 h with PMA/ionomycin. Restimulations were performed in the respective fed and starved media, and restimulation was initiated after the 24 h starvation period. (F, G) Cells were fed in full medium, or starved of cysteine for 24 h, then replated and co-cultured for 6 h with EL4-OVA lymphoma cells, in either fed or cysteine-starved media, at a ratio of 5:1 effector:target cells. EL-4 OVA cells had been pre-stained with CellTrace Violet. The percentage of dead EL4-OVA cells was measured as a readout of T cell cytotoxicity. (H, I) Proliferation and cell cycle analysis of fed and cysteine-starved cells by EdU/FxCycle Violet staining.

We also compared bioenergetics of effector T cells with in vitro-generated memory T cells^10^. SAA and glutamine starvation strongly blocked oxygen consumption rate (OCR), in effector T cells, with methionine and cysteine starvation also inhibiting OCR. However, OCR in memory T cells was unaffected by sulfur starvation (Figure S1G, S1H), indicating that sulfur is more important for rapidly proliferating, metabolically active cells, than for more quiescent memory cells with lower metabolic demands. ATP production was unaffected in all starvation conditions after 6 h (Figure S1I). 24 h cysteine starvation altered the surface marker profile on CD8^+^ T cells, decreasing CD44 expression while leaving CD62L unaffected, and significantly enhancing expression of the early activation marker CD69 (Figure S1J). Together, these results indicate that cysteine starvation enhances CD8^+^ T cell effector function, but decreases proliferation. These findings were specific to cysteine starvation, as 24 h methionine starvation decreased viability. Thus, we focused on exactly how cysteine is utilized by CD8^+^ T cells.

### CD8^+^ T cells acquire cysteine and use it to make glutathione

We incubated activated CD8^+^ T cells with stable isotope-labelled cysteine (^13^C,^15^N-Cys), extracted metabolites, and analyzed these extracts by nuclear magnetic resonance (NMR) (Figure 2A). ^13^C-^1^H 2D NMR peaks representing the α and β carbons of ^13^C,^15^N-Cys were detected in extracts fromcells incubated with ^13^C,^15^N-Cys (Figure 2A(i)). Extracts from cells incubated with ^13^C,^15^N-Cys had ^13^C-^1^H peaks representing the α (box #) and β (box *) carbons of the cysteine component of the glutathione (GSH) tripeptide, indicating that acquired labeled cysteine is metabolized into GSH (Figure 2A(i)). Confirming this, the peaks representing the GSH carbons were abolished upon treatment with the GSH synthesis inhibitor buthionine sulfoximine (BSO) (Figure 2A(ii)). Peaks for GSH and cysteine carbons were absent in cysteine-starved cells (Figure 2A(iii)), confirming that omitting cysteine from media results in intracellular cysteine depletion. We verified peak identities by acquiring NMR spectra for cysteine and GSH standards (Figure S2). We confirmed these results by LC-MS, detecting ^13^C from ^13^C-cysteine (Figure S3A, B) in reduced (GSH) and oxidized (GSSG) glutathione, as well as the GSH metabolite cysteinylglycine (CysGly) (Figure 2B). ^13^C was also detected in acetyl-coenzyme A, which contains the cysteine-derived cofactor coenzyme A (CoA), but is made in a separate pathway to GSH. We directly measured glutathione content using a luminescence assay, and found that GSH and GSSG were depleted in cysteine-starved, but not methionine- or glutamine-starved cells, indicating that this effect was specific to cysteine (Figure 2C). Cysteine-starved cells also contained less cysteinylglycine, γ-glutamylcysteine, and acetyl-CoA than fed cells (Figure 2D).

**Figure 2.**
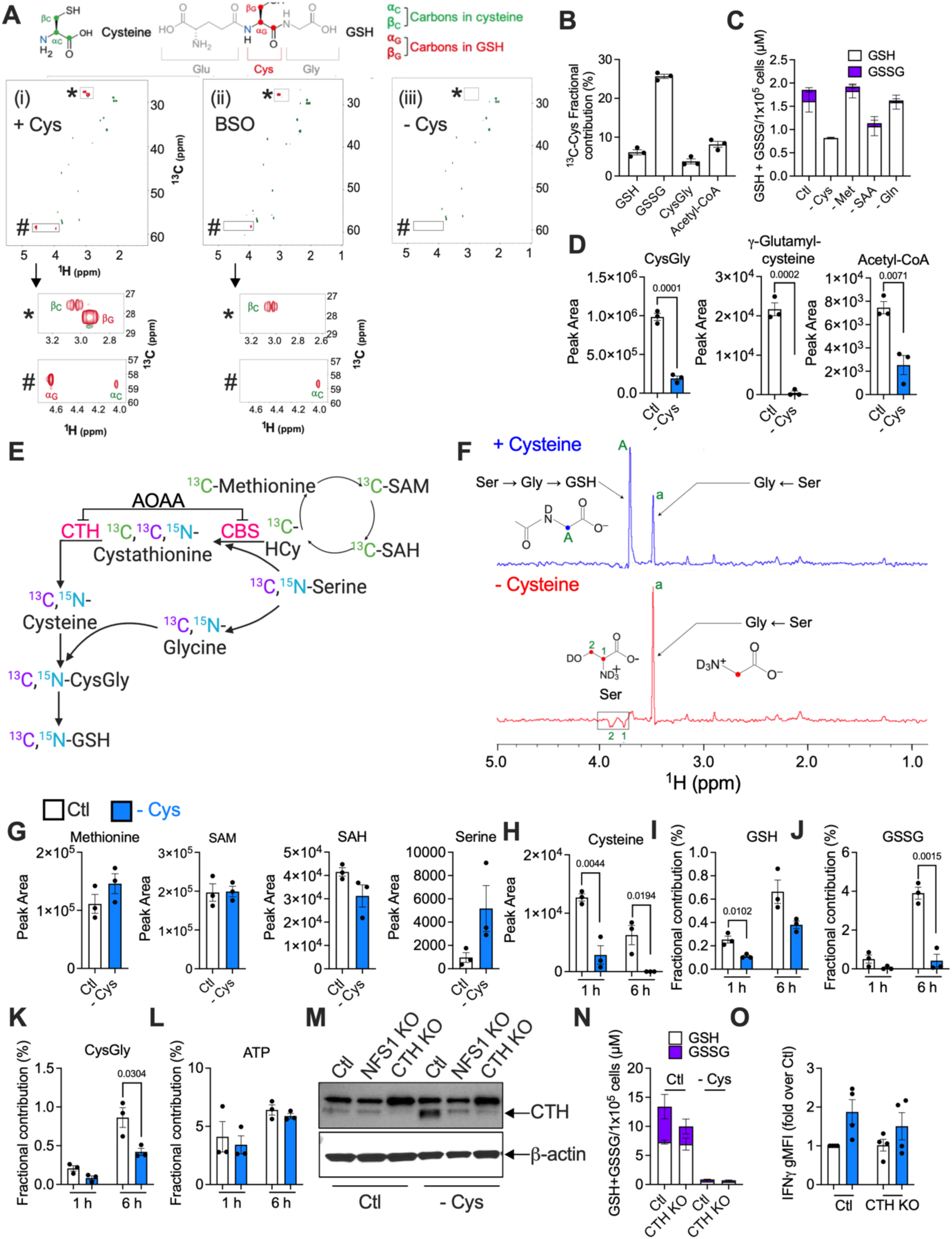
CD8^+^ T cells do not transsulfurate methionine to cysteine. (A) 40x10^6^ CD8^+^ T cells were fed with ^13^C-Cys (i), fed with ^13^C-Cys and treated with 0.5 mM BSO (ii) or starved of Cys (iii) for 6 h. 2D ^13^C-^1^H NMR analysis shows ^13^C-Cys uptake and metabolism to GSH. (B) Fractional contribution of ^13^C-Cys to metabolites, by LC-MS analysis. (C) Luminescence assay for GSH and GSSG in fed cells, or cells starved of the indicated amino acids (6 h). (D) LC-MS pool sizes of cysteine metabolites (24 h starvation). (E) Schematic indicating the isotope labeling strategy to probe methionine transsulfuration to cysteine by NMR. (F) NMR traces showing GSH, glycine, and serine peaks in T cells incubated with ^13^C,^15^N-Ser and ^13^C-Met. (G, H) LC-MS pool sizes of metabolites in fed or Cys-starved cells, after 24 h (G), or 1 and 6 h (H). (I - L) fractional contribution of ^13^C-Ser to GSH (I), GSSG (J), CysGly (K), and ATP, a known serine product (L) after 1 – 6 h. (M) Western blot showing the transsulfuration enzyme CTH in fed (Ctl) or cysteine-starved (- Cys, 24 h), wild type (Ctl) or CTH KO T cells. (N) Luminescence assay for GSH and GSSG and (O) IFNγ after PMA/ionomycin restimulation in fed or starved WT or CTH KO T cells.

### Activated CD8^+^ T cells do not transsulfurate methionine to cysteine

Many cells can transsulfurate methione into cysteine^11,12^, but there have been conflicting reports^13–17^ on whether this operates in T cells (Figure 2E). While methionine is an EAA, cysteine is considered non-essential on account of this transsulfuration pathway. We reasoned that if transsulfuration occurred in activated CD8^+^ T cells, methionine would replenish cysteine and, consequently, GSH. We concurrently traced ^13^C,^15^N-serine and ^13^C-methionine into activated CD8^+^ T cells, and analyzed metabolites by NMR (Figure 2E, F). In transsulfuration, methionine provides the sulfur atom for cysteine, while serine donates the carbon and nitrogen skeleton (Figure 2E). Therefore, if transsulfuration is engaged, ^13^C from methionine would be detected in the intermediate cystathionine, which then reacts with serine to form cysteine, and ^13^C and ^15^N from serine would be detected in both cystathionine and cysteine. If this cysteine is then used for GSH, ^13^C and ^15^N from serine would also be detected in cysteinylglycine and GSH/GSSG. Serine can also be directly metabolized to glycine, in which case ^13^C and ^15^N from serine will appear in glycine (and subsequently the *glycine* carbons and nitrogens in cysteinylglycine and GSH/GSSG). NMR indicates exactly which atom in a metabolite is labeled, i.e. it differentiates between GSH carbons that came from cysteine versus those from glycine. In fed cells (+ Cysteine), ^13^C from serine is detected in glycine (peak a), and in the glycine carbon of GSH (peak A) (Figure 2F, top panel). The labelled serine is not used to make cysteine, as no labelled cystathionine or cysteine are detected, and the cysteine carbon in GSH is unlabelled, and therefore cannot have come from serine. We questioned if transsulfuration might only become important when cysteine is limiting (- Cysteine). In this case, ^13^C from serine was still detected in glycine, and in residual serine (Figure 2F, bottom panel, inverse peaks 1 and 2). Again, cysteine was not labelled, and there was no GSH peak, indicating that it was not synthesized, even though the transsulfuration substrates were present (Figure 2G). Serine was confirmed to be elevated in cysteine-starved cells, by LC-MS (Figure 2G). Taken together, these results indicate that transsulfuration does not occur in these cells, and so serine and methionine cannot replenish cysteine and GSH.

To investigate when transsulfuration might occur during cysteine starvation, we traced ^13^C-methionine (Figure S3C) or ^13^C-serine in CD8^+^ T cells that had been fed or cysteine-starved for 1, 6, or 24 h, by LC-MS, which is more sensitive than NMR. Cysteine was progressively depleted from 1 – 6 h (Figure 2H), but methionine levels were similar in fed and cysteine-starved cells at 1, 6, and 24 h (Figure S3D, E). GSH, GSSG, and cysteinylglycine were abrogated in cysteine-starved cells after 24 h, with this depletion evident at 1 and 6 h (Figure S3D, F, G). The fractional contribution of ^13^C-serine to each of GSH, GSSG, and cysteinylglycine was lower at 1, 6 (Figure 2I - K), and 24 h (Figure S3H) in cysteine-starved cells compared to fed cells. Serine is also used for nucleotide synthesis; its fractional contribution to ATP was unaffected in cysteine-starved cells (Figure 2L), and ATP levels were maintained (Figure S3I), indicating that while glutathione synthesis was impaired, other serine metabolic pathways remained intact.

To genetically target transsulfuration we deleted cystathionine-γ-lyase (CTH) (Figure 2E) by CRISPR/Cas9 targeting, 2 days after activation with αCD3/28 + IL-2. CTH increased upon cysteine starvation and was successfully deleted (Figure 2M). We also silenced NFS1, a downstream cysteine desulfurase that utilizes available cysteine (for FeS cluster synthesis and thiolation reactions). NFS1 deletion had no effect on CTH levels in fed cells (Figure 2M). CTH can use GSH as a substrate to make reactive sulfur species and glutathionylate proteins to shield them from oxidative damage during redox stress^18–22^, possibly explaining its increase in cysteine-starved conditions. CTH deletion had no effect on GSH or GSSG levels in fed or cysteine-starved cells (Figure 2N), nor did it impact IFNγ or TNF production (Figure 2O, S3J). We tested CTH-deficient T cell activity in response to infection *in vivo*, a physiological nutrient environment. We transferred control or CTH-deficient CD45.2^+^ CD8^+^ OT-I T cells into CD45.1^+^ mice that had been infected one day prior with *Listeria monocytogenes*-expressing ovalbumin (LmOVA). 7 days post-transfer, control and CTH-deficient donor cells expanded to the same extent in infected mice (Figure S3K), expressed similar levels of the activation markers CD44, CD62L, and CD69 (Figure S3L), and produced comparable amounts of IFNγ and TNF upon PMA/ionomycin restimulation (Figure S3M). We also used aminooxyacetic acid (AOAA), a general pyridoxal phosphate (PLP)-dependent enzyme inhibitor that inhibits cystathionine-β-synthase (CBS) and CTH (Figure 2E), and other PLP-dependent enzymes^23^. AOAA increased S-adenosyl-homocysteine (SAH, Figure S3N), indicating CBS inhibition. Despite this, cysteine, cysteinylglycine, GSH, and GSSG levels were maintained in fed, AOAA-treated cells, and were not further lowered in AOAA-treated, cysteine-starved cells compared to starvation alone (Figure S3N, O). AOAA did not increase IFNγ in fed or cysteine-starved cells (Figure S3P). Together, these results demonstrate that activated CD8^+^ T cells do not engage transsulfuration of methionine to cysteine, making cysteine an EAA for these cells, in vitro and in a model of *Listeria* infection in vivo. We reasoned that if methionine transsulfuration replenished cysteine during cysteine starvation, deleting the transsulfuration machinery should exacerbate cysteine starvation to further decrease GSH, increase IFNγ, and inhibit proliferation, relative to cysteine starvation alone. Thus, the lack of any effect of CTH deletion further argues against a functional transsulfuration pathway in activated CD8^+^ T cells.

### Depleting GSH enhances cytokine production

We manipulated GSH in fed and cysteine-starved CD8^+^ T cells to examine its importance as a fate of cysteine. We confirmed that BSO (Figure S4A) depletes GSH, and that exogenous GSH recovers intracellular glutathione (Figure 3A). BSO did not affect proliferation in either fed or cysteine-starved cells (Figure 3B, C), and while GSH addition did not enhance proliferation in fed cells, it rescued the proliferative defect in cysteine-starved cells (Figure 3B). These data indicate that when cells have sufficient cysteine, GSH is not needed to maintain proliferation, however, GSH can support proliferation when cysteine is absent. Even though GSH is a major cysteine fate, its depletion does not mimic cysteine starvation in terms of its effect on proliferation. Surprisingly, BSO treatment, or CRISPR/Cas9 deletion of GCLC, the enzyme targeted by BSO (Figure S4A – C), enhanced IFNγ production (Figure 3D), indicating that continued GSH synthesis in fully activated T cells *restrains* IFNγ production, mimicking the effect of cysteine starvation on IFNγ. BSO-treated and GCLC-deficient T cells remained viable (Figure S4D). Even in fed cells, GSH supplementation decreased IFNγ, but not TNF (Figure S4E). It was important to acutely inhibit GCLC with BSO, or delete this enzyme after the first two days of activation, as GSH is required for the initial phase of CD8^+^ T cell activation and differentiation^24^. However, after that point, our results indicate that it becomes dispensable for T cells, and dampens effector function. Overall, these results indicated that cysteine starvation enhanced T cell function by abolishing GSH synthesis.

**Figure 3.**
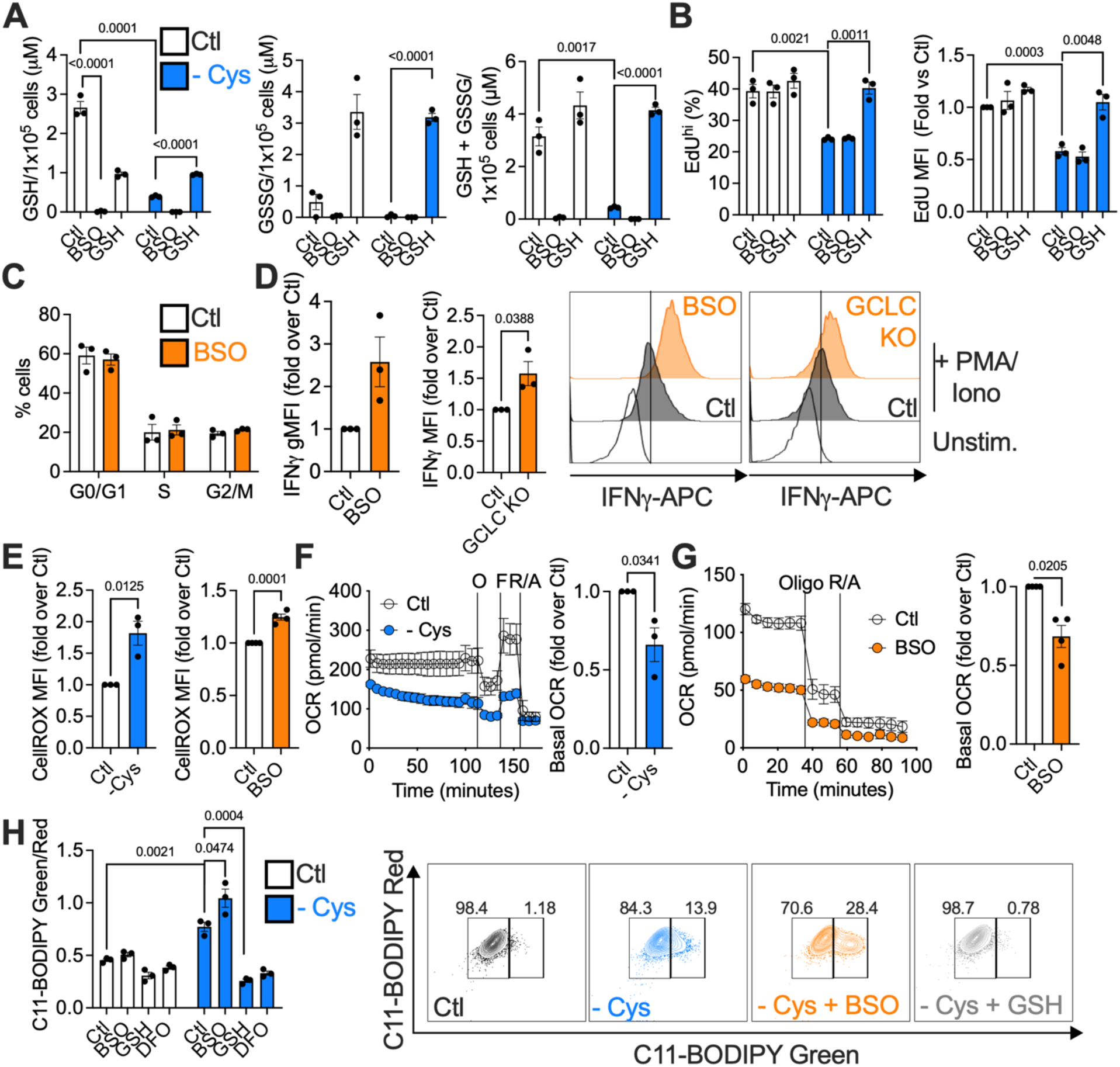
Depleting GSH enhances cytokine production while maintaining proliferation. T cells were fed or cysteine-starved (- Cys) for 24 h, and were concurrently treated with 0.5 mM BSO, 1 mM GSH, or left untreated. (A) GSH, GSSG, and total glutathione by luminescence assay. (B) Proliferation as measured by EdU incorporation. (C) Cell cycle analysis by EdU/FxCycle Violet staining in fed, BSO-treated cells. (D) IFNγ production after PMA/ionomycin restimulation of fed, BSO-treated, WT or GCLC-deficient (GCLC KO) T cells. (E) ROS production by CellRox staining of cys-starved (left) or BSO-treated (right) cells. (F, G) OCR of cys-starved (F) or BSO-treated (G) T cells by Seahorse analysis. (H) C11-BODIPY staining for lipid peroxidation. DFO was used as a negative control for lipid peroxidation.

We wondered if inhibiting GSH synthesis caused unsustainable oxidative damage in activated CD8^+^ T cells. Cysteine starvation or BSO treatment increased reactive oxygen species (ROS) (Figure 3E), and lowered OCR (Figure 3F, G), indicating decreased mitochondrial respiration. Like BSO-treated cells, GCLC KO T cells had increased ROS (Figure S4F), but neither BSO nor GCLC deletion affected mitochondrial mass (Figure S4G). Another redox pair, NAD^+^/NADH, was unaffected by cysteine starvation (Figure S4H), implying that redox balance was not globally impacted, and was restricted to cysteine-dependent GSH/GSSG. Nuclear factor erythroid 2-related factor 2 (NRF2), a major transcription factor regulating responses to oxidative damage, and its target, heme oxygenase-1 (HO-1), were increased in cysteine-starved cells, but were restored to normal levels by GSH supplementation (Figure S4I, J). Importantly, BSO treatment did not induce lipid peroxidation in fed cells, but exacerbated the increased lipid peroxidation caused by cysteine starvation (Figure 3H). This finding revealed that GSH was not necessary to protect against oxidative damage, as long as cysteine was present. Having neither cysteine nor GSH became problematic, greatly increasing lipid peroxidation. This begged the question of what cysteine does to protect against oxidative damage and promote cell function, other than making GSH. Of note, we found that the iron chelator deferoxamine (DFO) prevented lipid peroxidation during cysteine starvation (Figure 3H), indicating that cysteine starvation drives such peroxidation by increasing free iron. Via the Fenton reaction, Fe^2+^ promotes generation of the oxygen radicals that cause lipid peroxidation and cell death in ferroptosis^25,26^. Thus, these results suggest that cysteine may protect against oxidative damage by regulating iron.

### NFS1 supplies sulfur from cysteine for FeS cluster synthesis to support CD8^+^ T cell activity

The fact that GSH was not necessary to support T cell proliferation and counter oxidative damage in a cysteine-replete cell implied that an alternate fate of cysteine mediated these effects. The cysteine desulfurase NFS1 is a central regulator of sulfur metabolism from cysteine. It extracts the sulfur atom from cysteine (and not methionine) to make new FeS clusters and enact thiolation reactions. FeS clusters are critical cofactors for many enzymes, including components of the electron transport chain (ETC), and enzymes controlling ribosome recycling, translation, DNA replication, iron handling, and proliferation^8,9^. Given that the iron chelator DFO blocked T cell lipid peroxidation upon cysteine starvation, we questioned if cysteine supported T cell proliferation by fueling NFS1 as a central regulator of iron-sulfur metabolism.

We first tested if intact NFS1 function was needed to support T cell activity. Control or NFS1-deleted naïve (day 0 of culture) CD45.2^+^ CD8^+^ OT-I T cells were adoptively transferred into CD45.1^+^ recipient mice that had been infected one day prior with LmOVA. Control donor cells expanded in response to infection, while NFS1-deficient T cells failed to proliferate (Figure 4A, S5A). To investigate the role of NFS1 in activated CD8^+^ T cells in vitro, we deleted NFS1 by CRISPR/Cas9 after two days of T cell activation (Figure 4B). NFS1-deficient cells maintained CD44 and CD62L expression similarly to controls (Figure S5B). NFS1 deletion decreased expression of FeS cluster-containing enzymes, such as SDHB and NDUFB8, complexes I and II (CI, CII) of the ETC, respectively, as well as ISCU, part of the FeS synthesis machinery (Figure 4B), and cytosolic thiouridylase subunit 2 (CTU2), an enzyme that interacts with NFS1 to thiolate tRNA (Figure S5C). NFS1 silencing also lowered levels of a non-mitochondrial FeS-containing enzyme, aconitase, but did not affect TOMM20, a mitochondrial protein that does not have a FeS cluster (Figure S5D, E), or mitochondrial mass (Figure S5F). Agreeing with the reduced expression of ETC components, OCR was lowered, indicating decreased mitochondrial respiration (Figure 4C). NFS1-silencing mirrored the effect of cysteine starvation on proliferation. The percentage of highly proliferative (EdU^+^ FxCycle Violet^+^) cells was lower in the NFS1-deficient population, and these cells were stalled in the G0/G1 phase of the cell cycle (Figure 4D, E). NFS1-deficient T cells produced less IFNγ (Figure 4F), but more TNF (Figure 4G) than control T cells. Thus, NFS1 deficiency did not phenocopy cysteine starvation in terms of IFNγ production, indicating that continued NFS1-dependent cysteine metabolism was needed for IFNγ. We questioned if increased GSH production in the absence of NFS1 activity dampened IFNγ, as cysteine supply is not limiting, and cysteine could still be used to make GSH. Indeed, NFS1-deficient T cells produced more GSH, GSSG, and the GSH synthesis intermediate γ-cysteinylglycine, than control cells (Figure 4H, S5G), and BSO treatment was still capable of increasing IFNγ production in NFS1 KO cells (Figure 4I), reinforcing the idea that GSH levels regulate IFNγ production. NFS1-deficient T cells also had less ROS (Figure S5H), indicating enhanced antioxidant capacity, and NFS1 deficiency, unlike cysteine starvation, did not induce NRF2 or HO-1 expression (Figure S5I, J).

**Figure 4.**
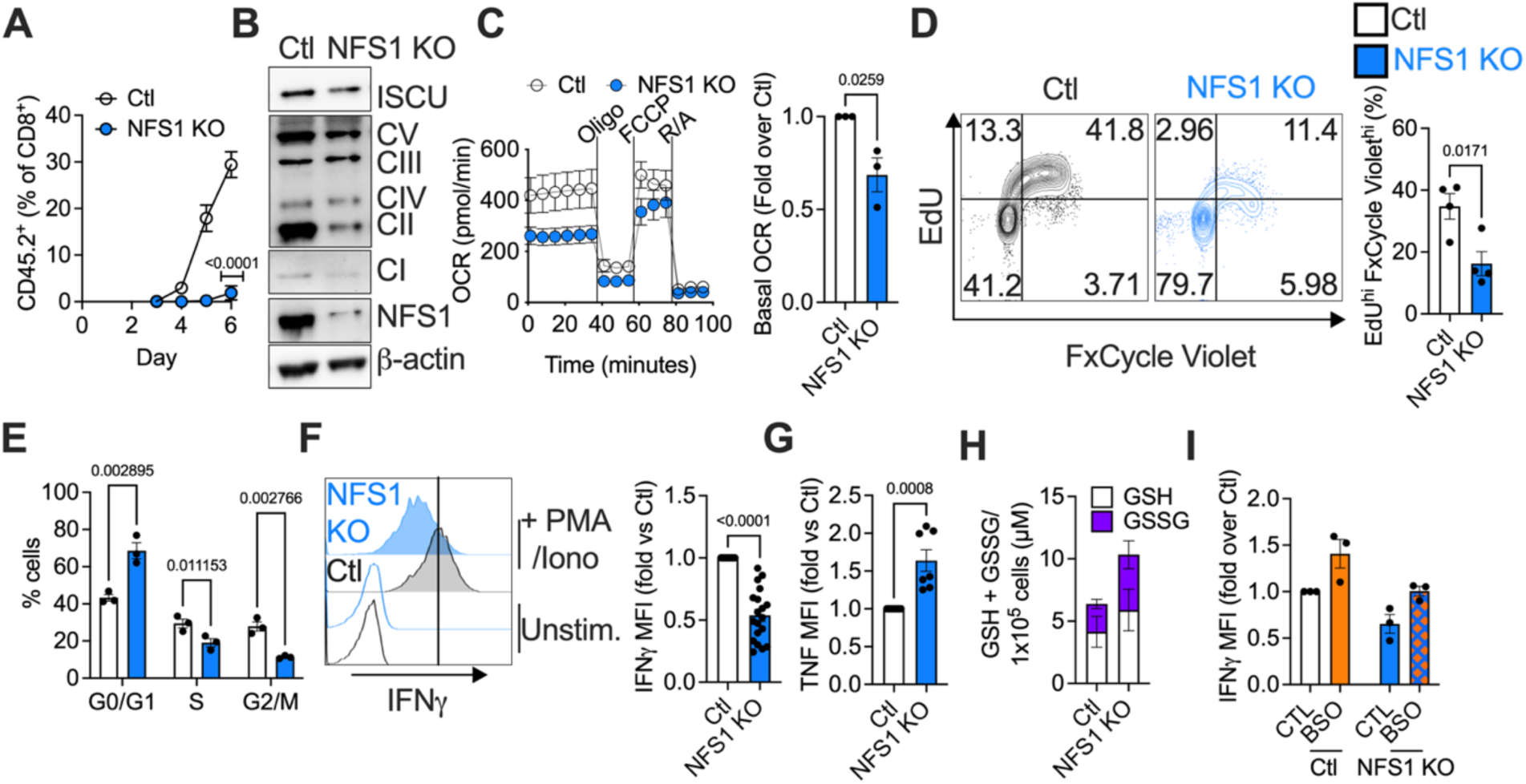
Sulfur metabolism via NFS1 supports proliferation and cytokine production. (A) NFS1 was deleted by CRISPR/Cas9 targeting in isolated naïve CD8^+^ T cells (day 0), and 0.1x10^6^ control or NFS1-deficient OT-I CD45.2^+^ CD8^+^ T cells were transferred into CD45.1^+^ mice that had been infected one day previously with 5x10^6^ LmOVA. Mice were bled daily and expansion of CD45.2^+^ CD8^+^ donor T cells was measured. (B - I) NFS1 was deleted by CRISPR/Cas9 in CD8^+^ T cells after 2 days of activation. (B) Western blot showing NFS1 deletion, and levels of ISCU, and OXPHOS proteins. (C) Seahorse analysis of OCR. (D, E) EdU/FxCycle Violet staining measuring proliferation and cell cycle stages. (F, G) IFNγ (F) and TNF (G) levels in cells that had been restimulated for 5 h with PMA/ionomycin. (H) GSH and GSSG levels by luminescence assay. (I) IFNγ in cells that had been treated with 0.5 mM BSO for 24 h prior to restimulation with PMA/ionomycin.

To investigate if GSH could still rescue defective proliferation due to cysteine starvation in a NFS1-deficient T cell, we starved WT or NFS1 KO cells of cysteine, +/- GSH supplementation. As before, cysteine starvation elevated IFNγ in WT cells, which was reversed by GSH, to levels lower than in WT, fed cells (Figure 5A). We reasoned that supplementing WT, cysteine-starved cells with GSH should mimic NFS1 deficiency, as the GSH arm of sulfur metabolism is preserved, but the substrate for NFS1 is absent, thereby abolishing NFS1 activity. Supporting this idea, T cells deleted for NFS1 produced the same levels of IFNγ as WT, starved cells supplemented with GSH (Figure 5A). Cysteine starvation or NFS1 deletion reduced proliferation, with these perturbations having an additive effect. GSH restored proliferation in a WT, cysteine-starved cell, but could not overcome NFS1 deficiency, only recovering proliferation to the level of a fed, NFS1-deficient cell, rather than a fed, WT cell (Figure 5B, C). These results implied that GSH supports proliferation independently of NFS1.

**Figure 5.**
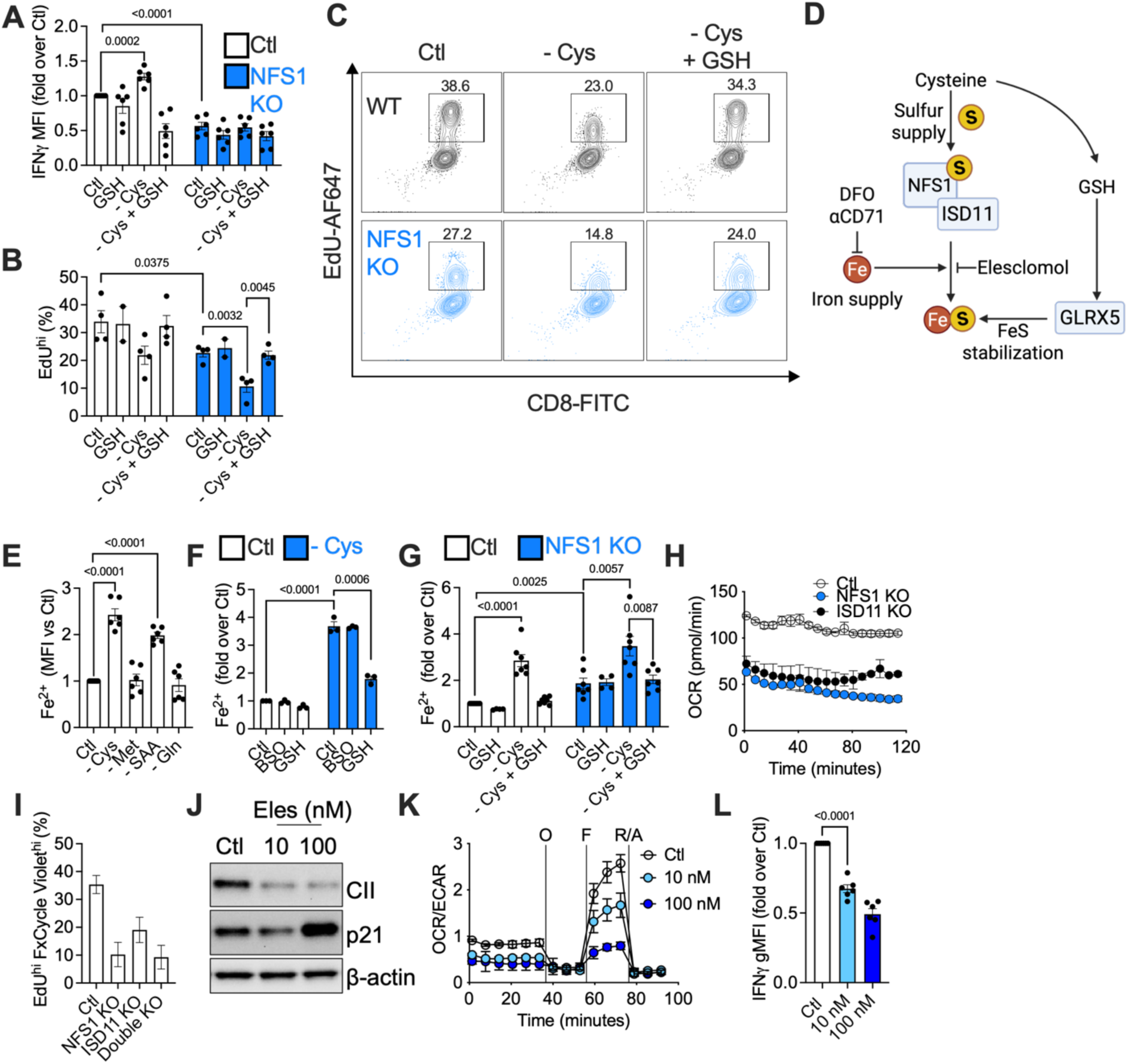
GSH and NFS1 support proliferation and cytokine production via FeS cluster stabilization and synthesis. (A – C) Control (Ctl) or NFS1 KO T cells were fed or cysteine-starved (24 h), +/- GSH (1 mM, 24 h, concurrent with cysteine starvation). (A) IFNγ in cells that were restimulated for 5 h with PMA/ionomycin. (B, C) EdU staining measuring proliferation. (D) Schematic diagram describing FeS cluster synthesis. (E - G) Fe^2+^ staining using BioTracker far-red labile Fe^2+^ dye. (E) Cells were starved of the indicated amino acids for 6 h. (F) Cells were fed or cysteine-starved, and treated with BSO (0.5 mM) or GSH (1 mM) concurrent with the 24 h cysteine starvation. (G) Control (Ctl) or NFS1 KO T cells were fed or cysteine-starved (24 h), +/-GSH (1 mM, 24 h, concurrent with cysteine starvation). (H) OCR by Seahorse analysis of Ctl, NFS1 KO and ISD11 KO cells. (I) EdU/FxCycle Violet staining measuring proliferation. (J - L) Cells were treated with elesclomol (Eles; 10 – 100 nM) for 24 h. (J) Western blot measuring CII, the cell cycle inhibitor p21, and β-actin. (K) OCR/ECAR ratio, by Seahorse. (L) IFNγ production.

### Cysteine regulates FeS clusters, via GSH and NFS1, to support T cell proliferation and cytokine production

We asked how GSH and NFS1 might converge to regulate proliferation and cytokine production. NFS1 supplies sulfur atoms from cysteine for FeS synthesis, but FeS clusters can also be directly stabilized by GSH, independently of NFS1^27^ (Figure 5D). Considering our proliferation results (Figure 5B, C), NFS1-deficient T cells would not be able to make new FeS clusters, but existing clusters could be stabilized by GSH (and any still bound to NFS1 in the iron-sulfur cluster synthesis complex, ISC). In the case of cysteine-starvation, NFS1 KO cells would have neither new FeS clusters, nor would they be able to stabilize existing clusters, leading to the lowest levels of proliferation, but GSH supplementation would restore this stabilization. This would occur independently of NFS1, but would not solve the problem of NFS1 deficiency, so no new FeS clusters would be made, and proliferation would be at the level of a fed, NFS1 KO cell.

To investigate the synthesis of new FeS clusters in CD8^+^ T cells, we first measured free Fe^2+^, as an indicator of disrupted FeS formation. Intracellular Fe^2+^ could increase as a result of insufficient sulfur to complex it into FeS clusters, and also due to activation of the iron starvation response during FeS cluster disruption, which enhances iron import via the transferrin receptor (CD71)^28^. We used a dye to detect Fe^2+^, confirming its efficacy by loading our cells with iron using ferrous ammonium sulfate (FAS) or chelating iron using bipyridyl (BIP) (Figure S6A). Cysteine starvation specifically increased Fe^2+^; neither methionine nor glutamine starvation had the same effect (Figure 5E). BSO had no effect on Fe^2+^ in fed or cysteine-starved cells, but GSH supplementation reversed the increase in Fe^2+^ caused by cysteine starvation (Figure 5F), indicating that GSH modulates iron metabolism. NFS1 deletion also elevated Fe^2+^, which was exacerbated by cysteine starvation, and in turn, recovered (but only to the level of a fed, NFS1 KO cell) by GSH (Figure 5G). These changes in Fe^2+^ reflect the differences in proliferation observed in fed, starved, and GSH-supplemented WT and NFS1 KO T cells (Figure 5B, C), supporting our idea that disrupted FeS synthesis underlies these proliferative defects. The iron chelator DFO had no effect on GSH, GSSG, or proliferation in fed or cysteine-starved cells (Figure S6B, C), suggesting that disrupting iron metabolism does not initiate these observed changes in proliferation and glutathione, and altered iron metabolism is rather a consequence of perturbed sulfur metabolism. GSH rescued SDHB deficiency due to cysteine starvation in control cells, and this was also unaffected by DFO (Figure S6D). DFO did antagonize cysteine starvation-induced lipid peroxidation (Figure S6E), further indicating that iron accumulation occurred downstream of sulfur starvation and disrupted FeS cluster synthesis (and so could then drive lipid peroxidation), and was not an upstream event.

To investigate the source of the increased iron, we examined the iron transporter CD71 (transferrin). In agreement with induction of the established FeS cluster disruption response^28^, NFS1 deletion increased CD71 levels (Figure S6F). A blocking antibody against CD71 (αCD71) decreased intracellular Fe^2+^ in cysteine-starved wild type and NFS1-deficient T cells (Figure S6G, left), indicating that elevated iron import via CD71 contributed to the increased free iron observed due to cysteine starvation, NFS1 deletion, or both. However, cysteine starvation and NFS1 deletion still boosted Fe^2+^ in cells in which CD71 was blocked, with these perturbations having an additive effect (Figure S6G, right), indicating that part of the increase in free iron originates from disrupted FeS clusters within the cell, or as a result of insufficient sulfur to complex this iron.

Since NFS1 also has a role in tRNA thiolation, we more precisely probed the role of FeS synthesis by genetically targeting ISD11, the specific factor that complexes with NFS1 to make FeS clusters^29,30^ (Figure 5D). ISD11 deletion (Figure S6H) mimics NFS1 deficiency, increasing Fe^2+^ (Figure S6I), lowering OCR (Figure 5H, S6J), and proliferation (Figure 5I). Similar to NFS1 KO cells, ISD11 deletion arrests cells in G0/G1, leading to a lower fraction in the S and G2/M phases of the cell cycle (Figure S6K). Linking FeS cluster status to changes in proliferation, we noted that cysteine starvation or NFS1 deletion decreased levels of the ribosome recycling factor and translation initiation factor ABCE1, which is an FeS-dependent enzyme, while increasing expression of the cell cycle inhibitor p21 (Figure S6L). Elesclomol, a pharmacological inhibitor of FeS synthesis^31^ (Figure 5D), dose-dependently lowered levels of the FeS-containing enzyme CII, while increasing p21 (Figure 5J). This drug decreased the OCR/ECAR ratio, both reducing OCR and increasing ECAR (Figure 5K, S6M). Furthermore, elesclomol inhibited IFNγ production, with a more minor effect on TNF (Figure 5L, S6N). Collectively, these data show that directly inhibiting FeS synthesis mimics NFS1 deletion, blocking proliferation and IFNγ production.

We probed how GSH might stabilize FeS clusters independently of NFS1. Glutaredoxins (GLRXs) 1, 2, 3, and 5 use GSH as a cofactor to regulate redox balance, for example by recycling thioredoxins and reducing disulfides (GLRX1, 2), and stabilizing cytosolic (GLRX3) and mitochondrial (GLRX5) FeS clusters^32^. CRISPR/Cas9 targeted deletion of GLRX5, but not of GLRX1/2/3, decreased CII levels (Figure S6O), and CII could not be rescued by GSH supplementation (Figure S6P), indicating that GLRX5 may be one mechanism by which GSH regulates FeS clusters independently of NFS1 (Figure 5D). GLRX5 deletion increased Fe^2+^, impeded proliferation of adoptively transferred CD8^+^ T cells in response to LmOVA in vivo, and dampened IFNγ and TNF production in vitro (Figure S6Q – S), indicating that disrupting GSH/GLRX5-mediated FeS cluster stabilization impacts CD8^+^ T cell proliferation and function in a similar manner to blocking new FeS cluster synthesis.

### GSH is not catabolized to replenish cysteine

We considered the possibility that besides directly stabilizing FeS clusters, GSH was itself broken down to replenish cysteine. Two intracellular enzymes, glutathione-specific γ-glutamylcyclotransferases (CHACs) 1 and 2^33–35^, and an ecto-enzyme, γ-glutamyltransferase 1 (GGT1) can catabolize GSH (Figure S7A) into cysteinylglycine, which can be further broken down into cysteine and glycine, and a γ-glutamyl-amino acid, which is then metabolized to 5-oxoproline. CHAC1 can be induced in response to cystine starvation in cancer cell lines^36^, restoring cysteine to supply sulfur for FeS clusters. In the case of GGT1, cyste(i)ne is generated extracellularly, and then acquired by the cell. CHAC1 is not expressed in either fed or cysteine-starved CD8^+^ T cells (Figure S7B). CHAC2 is constitutively expressed in these cells, and we knocked it out by CRISPR/Cas9 targeting (Figure S7C). Neither cysteine starvation nor GSH supplementation modulated CHAC2 levels, and GSH supplementation still restored SDHB (CII) expression in cysteine-starved, CHAC2-deficient T cells (Figure S7C). We expected that if CHAC2 catabolized GSH to boost cysteine, then in its absence GSH would accumulate, but GSH and GSSG levels were the same in wild type and CHAC2 KO T cells, in fed, cysteine-starved, and GSH-supplemented conditions (Figure S7D). Similarly, CHAC2 deficiency had no impact on proliferation, or IFNγ or TNF production (Figure S7E – G). These results argue that CHAC2 does not catabolize GSH to restore cysteine in our system.

We also investigated if GGT1 could break down extracellular GSH to replenish cysteine^37,38^. 6 h cysteine, methionine, or glutamine starvation had no effect on GGT1 activity, as measured by production of the fluorescent substrate AMC in a GGT1 activity assay (Figure S7H), but 24 h cysteine starvation increased AMC production, indicating increased GGT1 activity (Figure S7I). The pharmacological GGT1 inhibitor GGStop^39^ successfully blocked GGT1 activity, in fed, cysteine-starved, and GSH-supplemented cells (Figure S7J, K). However, GGStop had no effect on GSH-mediated rescue of SDHB or ABCE1 expression in cysteine-starved cells, nor did it exacerbate the proliferative defect due to cysteine starvation, as measured by p21 levels (Figure S7L). There were no differences in IFNγ or TNF in the presence of GGStop (Figure S7M, N). Together, these findings indicate that inhibiting GGT1 activity did not compound the effects of cysteine starvation, nor did it interfere with the ability of GSH to rescue these defects, suggesting that GGT1 does not restore cysteine levels in these cells.

### NFS1-dependent cysteine metabolism enhances anti-tumor immunity

Given the importance of NFS1-dependent cysteine metabolism supporting proliferation and cytokine production in CD8^+^ T cells, we examined the relevance of this pathway in an in vivo tumor model. We injected B16-OVA melanoma cells into the flanks of CD45.1^+^ recipient mice and, once tumors were palpable, adoptively transferred control or NFS1-deficient CD45.2^+^ CD8^+^ OTI T cells into the tumor-bearing mice. Tumors grew larger in mice receiving NFS1-deficient T cells (Figure 6A – C, Figure S8A), indicating that NFS1 deficiency impairs CD8^+^ T cell anti-tumor immunity. Infiltration of adoptively transferred NFS1 KO CD8^+^ T cells into the tumor was impaired (Figure 6D,E), and NFS1-deficient cells had increased expression of the exhaustion marker TIM3 (Figure 6F). LC-MS analysis of tumor interstitial fluid (ISF) showed that cysteine was elevated in the tumor microenvironment (TME) in mice that had received NFS1-deficient T cells compared to those that had received control T cells (Figure 6G), indicating that changes in nutrient utilization by TILs can shape the TME. This was likely not just a consequence of decreased proliferation of NFS1 KO T cells, as other metabolites, including tyrosine (Figure 6H), glutamine, and methionine (Figure S8B) were unaffected, i.e. TME metabolite levels were not globally increased in mice that had received NFS1 KO T cells.

**Figure 6.**
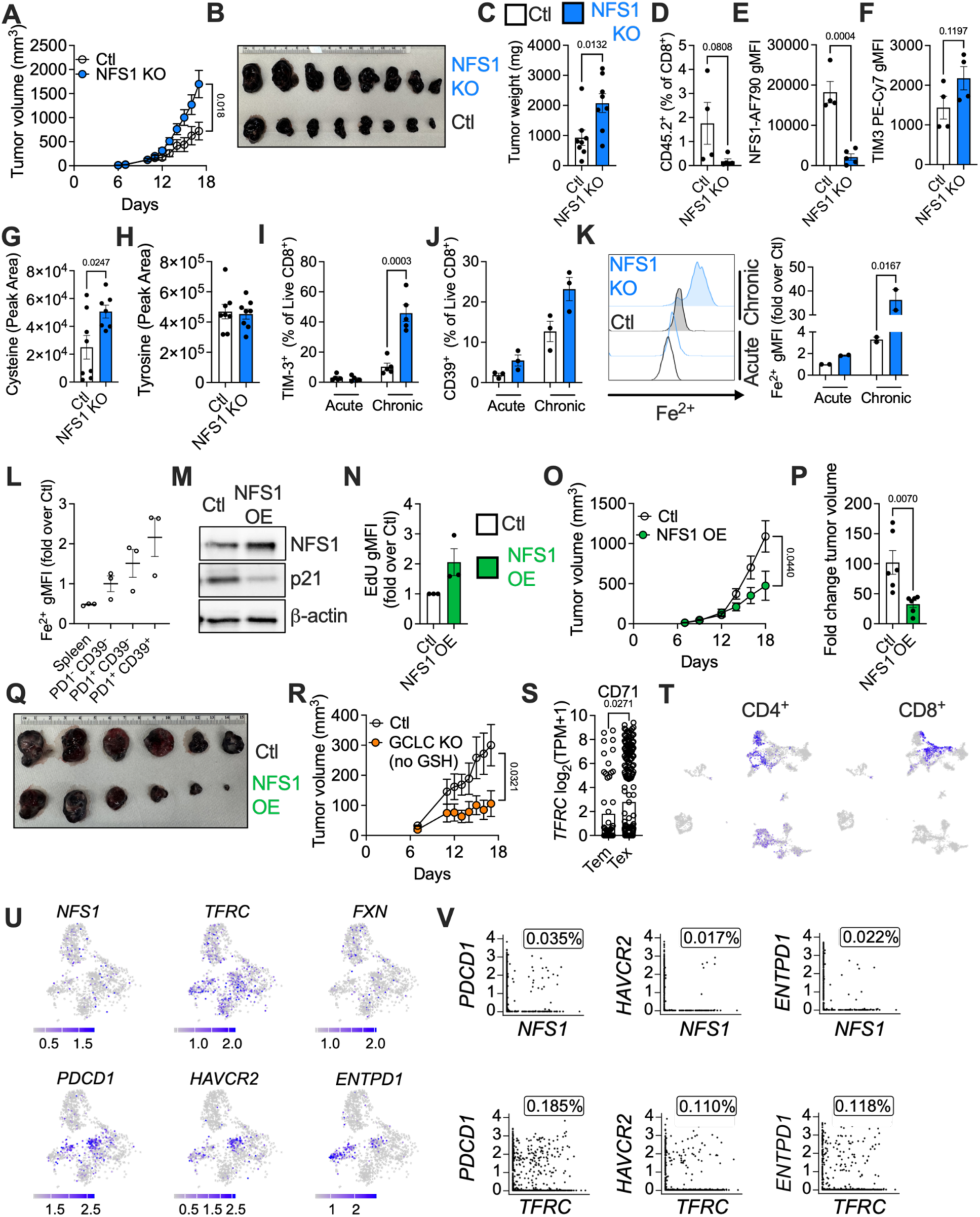
NFS1 metabolism promotes CD8^+^ T cell function in vivo. (A - H) 1x10^6^ B16-OVA melanoma cells were subcutaneously injected into the flanks of CD45.1^+^ recipient mice. 7 days post-implantation, when tumors were palpable, 0.5x10^6^ control (Ctl) or NFS1 KO CD45.2^+^ CD8^+^ OT-I T cells were adoptively transferred into these recipients by intravenous tail vein injection. Tumor growth curve (A), final tumor size (B), weight (C), and donor T cell infiltration (D), NFS1 (E), and TIM3 (F) expression were measured. (G, H) metabolites in tumor institial fluid were measured by LC-MS. (I - K) In an in vitro model of T cell exhaustion, control or NFS1 KO cells were initially activated for 2 days with αCD3/28 + IL-2, then cultured with IL-2 only (Acute) or restimulated with IL-2 + αCD3/28 (Chronic) every 48 h for 8 days. The frequency of TIM-3^+^ (I) and CD39^+^ (J) CD8^+^ T cells was measured, as well as Fe^2+^ levels (K). (L) CD8^+^ T cells from spleen, and tumor-infiltrating lymphocytes (TILs) from B16 melanoma tumors, from the same mouse, were stained for exhaustion markers PD1, CD39, and Fe^2+^. (M) (O - R) B16-OVA melanoma tumors were established as in (A). (O – Q) 0.5x10^6^ control (Ctl) or NFS OE CD45.2^+^ CD8^+^ OT-I T cells were adoptively transferred into CD45.1^+^ recipients, and tumor growth (O), fold change in tumor size vs day 7 (P), and final tumor size (Q) were measured. (R) 0.5x10^6^ control (Ctl) or GCLC KO CD45.2^+^ CD8^+^ OT-I T cells were adoptively transferred into CD45.1^+^ recipients, and tumor growth was measured. (S) CD71 expression in effector memory (Tem) and exhausted (Tex) TILs in human HCC patients, from a previously published scRNAseq dataset. (T – V) A previously published human HCC scRNAseq analysis was reanalyzed. (T) CD4^+^ and CD8^+^ T cell clusters were identified. (U, V) CD4^+^ and CD8^+^ T cells expressing *NFS1*, *TFRC* (CD71), *FXN* (frataxin), and the exhaustion markers *PDCD1* (PD-1), *HAVCR2* (TIM-3), and *ENTPD1* (CD39) were identified (U), and co-expression of *NFS1* (top) or *TFRC* (bottom) with each of these exhaustion markers was determined (V).

To further investigate a role for NFS1 in T cell exhaustion suggested by our in vivo data, we used an in vitro model of T cell exhaustion by repetitive αCD3/28 restimulation^40,41^. We activated CD8^+^ T cells with αCD3/28 + IL-2 for 48 h, then maintained these cells in IL-2 (Acute) or IL-2 + αCD3/28 (Chronic) every 2 days for 8 – 10 days. NFS1-deficient cells exhibited accelerated exhaustion, expressing TIM3 (Figure 6I) and CD39 (Figure 6J) earlier than control cells. The frequency of PD1^+^ cells also increased more quickly in NFS1 KO compared to control cells, reaching its peak earlier, and was higher in acutely stimulated NFS1-deficient cells compared to controls, even without chronic stimulation (Figure S8C). Furthermore, this correlated with disrupted iron metabolism. Acute NFS1 KO T cells had higher Fe^2+^ than control cells, as previously observed (Figure 5G, 6K). Chronic control cells had increased Fe^2+^ relative to Acute control cells, associating Fe^2+^ disruption with exhaustion in the WT setting, and this was greatly exacerbated in Chronic NFS1-deficient cells, correlating with progression to exhaustion (Figure 6K). We measured Fe^2+^ in CD8^+^ tumor-infiltrating lymphocytes (TILs) from B16-OVA melanoma tumors, and similarly found that as CD8^+^ T cells acquire exhaustion markers (PD1^-^ CD39^-^ vs PD1^+^ CD39^-^vs PD1^+^ CD39^+^), intracellular Fe^2+^ progressively increased (Figure 6L). Redox balance was also disrupted in exhausted T cells (T_ex_): GSH levels remained steady, but GSSG levels increased during chronic stimulation (Figure S8D), indicating a more oxidative environment. Supporting this, cellular and mitochondrial ROS were increased in Chronic T cells, and were even further elevated in Chronic NFS1 KO cells, which had very low viability (Figure S8E – G). We also observed that NFS1 expression correlated with decreased exhaustion in a MC38 colon cancer model of immune checkpoint blockade (ICB) therapy. MC38 colon cancer cells were injected into the flanks of BALB/c mice, which were left untreated, or treated every second day from day 7 with ICB (α-PD-1 + α-CTLA4). ICB slowed tumor growth (Figure S8H). NFS1 expression was lower in CD8^+^, CD4^+^, and T_reg_ subsets from untreated mice compared to healthy spleens, but was recovered in all subsets by ICB treatment (Figure S8I - K).

To test if boosting NFS1 enhances CD8^+^ T cell function, we overexpressed NFS1 in CD8^+^ T cells (NFS1 OE) by retroviral transduction (Figure 6M, S8L, M). NFS1 OE T cells had increased proliferation, indicated by decreased p21 levels (Figure 6M) and higher EdU staining (Figure 6N). The FeS proteins SDHB (CII) and ABCE1 were also increased (Figure S8M). Adoptive transfer of NFS1 OE CD8^+^ OT-I T cells into mice bearing B16-OVA tumors improved tumor control compared to adoptive transfer of control CD8^+^ OT-I T cells. Tumors were smaller in mice that received NFS1 OE T cells (Figure 6O – Q, Figure S8N). In particular, the fold increase in tumor size was significantly decreased in mice that received NFS1 OE T cells, versus control cells (Figure 6P). Tumor sizes had been matched on the day of adoptive transfer (Figure S8O). Based on our in vitro data, we also questioned if diverting cysteine away from GSH synthesis improved effector function in vivo. We injected B16-OVA melanoma cells into the flanks of CD45.1^+^ recipient mice and, once tumors were palpable, adoptively transferred control or CD45.2^+^ CD8^+^ OTI T cells that had been deleted for GCLC two days after activation. GCLC-deficient T cells improved tumor control (Figure 6R), indicating that inhibiting GSH synthesis in fully activated CD8^+^ T cells enhances their anti-tumor immune function in vivo.

Analysis of a previously published single-cell RNAseq dataset^42^ characterizing TILs in human hepatocellular carcinoma (HCC) revealed that NFS1 and FeS metabolic pathways may be relevant for human anti-tumor immunity. Exhausted TILs expressing high levels of the exhaustion markers, TIM-3, CD39, and PD-1 (Figure S8P), compared to effector memory TILs, also had significantly elevated expression of CD71 (Figure 6S). We also analyzed a scRNAseq dataset^43^ of immune cells from multiple sites (liver tumors, adjacent liver, hepatic lymph nodes, ascites, and blood) in HCC patients (Figure S8Q), identifying clusters representing CD8^+^ and CD4^+^ T cells (Figure 6T). Within these clusters, cells expressing the exhaustion markers *PDCD1* (PD-1), *HAVCR2* (TIM-3), and *ENTPD1* (CD39) had low levels of *NFS1*, and of the FeS-synthesizing factor frataxin (*FXN*) (Figure 6U). However, T cells with high levels of *TFRC* (CD71) expression overlapped to a greater extent with exhausted T cells, mirroring our observations. The percentage of T cells co-expressing *NFS1* or *FXN* with any of these exhaustion markers was extremely low (Figure 6V, S8R), while a higher percentage (∼ 10-fold increase) of cells co-expressed *TFRC* and each exhaustion marker. Together, these findings suggest that decreased NFS1 and FeS synthesis, and consequently increased CD71 expression, may be associated with human T cell exhaustion in HCC, and that targeting NFS1, FeS clusters, or iron metabolism in human cancer may be of benefit.

## DISCUSSION

Activated CD8^+^ T cells augment cell surface expression of a subset of AA transporters^44,45^, acquiring specific AAs according to demand, instead of indiscriminately taking up all available AAs. EAAs must be acquired from dietary sources, while NEAAs can be synthesized to sufficient levels in the body. However, NEAAs can become conditionally essential when cellular demand outstrips their synthesis, and cells must then acquire NEAAs from external sources. Many physiological contexts in which NEAAs become conditionally essential remain to be defined, and here we show that the NEAA cysteine becomes conditionally essential for activated CD8^+^ T cells to proliferate and enact their effector function. We mechanistically demonstrate exactly how CD8^+^ T cells use cysteine, pinpointing metabolism of its sulfur atom as a key determinant of anti-tumor immunity.

Human CD8^+^ T cells increase cystine transport^46^ during an immune response, and T cell activation^16^, proliferation^17^, and DNA synthesis^47^ need cystine uptake. Various cancers show altered sulfur utilization^48–52^, for example, by increasing cysteine transport, decreasing its levels in the tumor microenvironment^6^, or by preserving sulfur metabolism, increasing methionine transsulfuration to cysteine^33^, or maintaining NFS1 activity^53^. Evidence for transsulfuration of methionine to cysteine in CD4^+^ T cells is mixed: activation of naïve primary human CD4^+^ T cells induced expression of cyste(i)ne transporters, and methionine was unable to compensate for cysteine starvation to maintain proliferation in these cells^17,47^, indicating that transsulfuration does not operate. However, work in human cell lines and primary murine CD4^+^ T cells^13,14^, tracing radiolabelled sulfur (^35^S) from methionine or homocysteine into GSH and cystine, respectively, suggests that transsulfuration may be possible here. This may be a feature of transformed cells, or CD4^+^ T cells may convert excess homocysteine to cyste(i)ne, but not engage the whole transsulfuration pathway from methionine (upstream of homocysteine) to cyste(i)ne and GSH. Our comprehensive NMR data demonstrate that transsulfuration of methionine to cysteine does not operate in activated CD8^+^ T cells, in contrast to many cancer cell types. This makes cysteine conditionally essential for CD8^+^ T cells, i.e. they must acquire it from their external environment to meet demands of increased proliferation, ROS buffering, and metabolic activity. This may serve the physiologic purposes of sparing methionine to provide methyl groups required for the nucleotide methylations and epigenetic reprogramming underlying T cell differentiation upon antigen stimulation^45^, and serine/glycine for nucleotide synthesis to support increased energy demands^54,55^. Methionine has recently been demonstrated to be rapidly consumed upon initial T cell activation, and is needed to maintain the SAM cycle and the intracellular methylome, regulating calcium signaling and protecting against exhaustion^56^. In this study, cysteine was also rapidly consumed upon T cell receptor (TCR) ligation. Overall, our results uncover a metabolic difference between CD8^+^ T cells and cancer cells, implicating transsulfuration inhibition as a potential anti-cancer target.

While GSH production is critical for the initial growth and metabolic reprogramming underlying T cell activation^24^, we reveal that its continued synthesis restrains effector function by activated CD8^+^ T cells. Surprisingly, inhibiting GCLC after the initial two days of activation enhanced IFNγ production, and improved anti-tumor immunity mediated by adoptively transferred CD8^+^ OT-I T cells in a B16-OVA melanoma tumor model. Exactly how GSH dampens cytokines remains an open question. Direct oxidative modifications control a vast array of enzyme activities relevant for protein synthesis, e.g. tRNA synthetases, which may alter cytokine production^57^. Transcription and translation factors, including eukaryotic initiation factor 2 (eIF2) and eIF4E binding protein 1 (4EBP1), are regulated by ROS-sensitive kinases such as PERK^58^ and mTORC1^59^. GSH itself can glutathionylate proteins to regulate their activity, including transcription and translation factors, and, intriguingly, GAPDH^60^, known to be a direct regulator of IFNγ^61^. Thus, there are a number of mechanisms by which GSH may control redox balance to influence cytokine production. GCLC blockade did not impair proliferation, or cause unsustainable oxidative damage and cell death, as long as there was still a supply of cysteine, suggesting that an alternate fate of cysteine maintained redox balance, proliferation, and cytokine production. A particularly telling finding was the fact that BSO exacerbated cysteine-induced lipid peroxidation, indicating that this was regulated by both GSH-dependent and -independent fates of cysteine. Further, the iron chelator DFO reversed starvation-induced peroxidation, suggesting that cysteine controlled redox balance by regulating intracellular iron.

Iron-sulfur chemistry arose early in evolution^62^; FeS proteins, and the enzymes that synthesize FeS clusters, are conserved from bacteria to humans^63^, but exactly how FeS biology controls immune cell function has not been investigated in detail, particularly from the sulfur angle. Previous work has examined iron import, metabolism, and ferroptosis as regulators of T cell immunity^64–69^, but sulfur is an important partner in many aspects of iron metabolism, especially at the level of FeS clusters, which acquire their sulfur component from cysteine. Critically, FeS clusters can also be stabilized by GSH, pinpointing FeS clusters as a point of control that can be regulated by cysteine, both via GSH and independently of GSH. Cysteine provides sulfur for newly synthesized FeS clusters via the cysteine desulfurase NFS1. NFS1 complexes with ISD11, acyl carrier protein (ACP), ferredoxin (FDX2), frataxin (FXN), and the scaffold protein ISCU to enact de novo FeS cluster synthesis in the mitochondria^70^, before new FeS clusters are guided to their target apoproteins in a chaperone-mediated process^71^. The utility of FeS clusters lies in their ability to transiently bind single electrons to facilitate electron transfer reactions, critical for intracellular processes such as the tricarboxylic acid cycle (aconitase)^72^, oxidative phosphorylation (ETC complexes)^8^, DNA replication^73^, and cell division^74^. Our results reveal that these fundamental cofactors support immune cell function: we show that activated CD8^+^ T cells use cysteine for new FeS cluster synthesis, via NFS1, in order to fully proliferate, and for protection against exhaustion, a major factor limiting immunity in chronic disease and cancer. NFS1-deficient T cells proliferate less and have increased expression of exhaustion markers, while enforcing NFS1 expression boosts levels of FeS proteins, enhances proliferation, and improves anti-tumor immunity in vivo.

Inhibited FeS cluster synthesis, due to either limited iron supply, or disruption of FeS-synthesizing enzymes, induces the iron starvation response, causing iron-responsive element binding proteins (IREs) to increase transferrin receptor (CD71) in an effort to acquire more iron^75,76^. Therefore, FeS clusters are a major determinant of intracellular iron and redox balance, regulating iron levels to prevent excessive iron-dependent ROS generation via Fenton chemistry. Our data show that T cells have multiple cysteine-dependent mechanisms to preserve FeS clusters, protecting against oxidative damage and ferroptosis. Synthesis of new FeS clusters via NFS1 prevents the iron starvation response, protecting against iron overload and maintaining redox balance. In the absence of NFS1, GSH achieves the same outcome by stabilizing existing clusters, e.g. via GLRX5, and by directly ligating free iron^77^. Only the total absence of GSH and NFS1 becomes intolerable for CD8^+^ T cells, abolishing proliferation and drastically elevating oxidative damage. This explains why after initial activation, CD8^+^ T cells can dispense with continued GSH synthesis, as long as they have cysteine – cysteine flux into FeS clusters via NFS1 is sufficient to protect against iron overload and maintain redox balance. Continued FeS cluster synthesis in our system may also regulate iron by stabilizing the FeS-dependent mitochondrial GSH importer Slc25A39, which has been described to import available GSH from the cytosol to the mitochondria during GSH depletion (for example, during BSO treatment), in HEK293T cells^78^. This GSH can then regulate free iron in the mitochondria. Disrupting FeS synthesis destabilizes Slc25a39, which leads to mitochondrial iron dysregulation. This phenomenon is in agreement with our results showing that the highest levels of free iron are observed in cells lacking both GSH and FeS synthesis.

Our findings held true in vivo. NFS1-deficient T cells, with impaired sulfur metabolism, exhibited poorer tumor control, while inhibiting cysteine flux into GSH (by deleting GCLC) or promoting NFS1-dependent sulfur metabolism improved anti-tumor immunity. Importantly, this may be relevant for T cell function in human cancers. Exhausted CD8^+^ T cells from human HCC patients have lower NFS1 expression, but increased CD71, compared to effector memory T cells, implicating iron-sulfur metabolism as a potential target to counter exhaustion and improve anti-cancer immune function in human patients. Overall, our detailed investigation of exactly how CD8^+^ T cells use cysteine provides a new level of metabolic control over cell function. Total nutrient restriction, either by starvation or transporter inhibition, would eliminate both beneficial and detrimental effects of that nutrient. Delineating exactly how different intracellular pathways use cysteine enables us to retain beneficial effects of cysteine while abolishing those that restrain function, at the appropriate times. We have illustrated this concept in the case of one metabolite, cysteine, but it is likely to apply to other metabolites relevant for immune cell function.

## ACKNOWLEDGMENTS

We thank members of the Pearce Labs for critical input during the project. This work was supported by NIH grant AI156274 and a Bloomberg Distinguished Professorship (to E.L.P.).

## MATERIALS AND METHODS

### Mice

C57BL/6J (RRID: IMSR_JAX:000664), major histocompatibility complex (MHC) class I-restricted OVA-specific TCR OT-I transgenic mice (RRID: IMSR_JAX:003831) and CD45.1^+^ C57/BL/6J (B6.SJL-*Ptprc^a^ Pepc^b^*/BoyJ;Jax, 002014) mouse strains were purchased from the Jackson Laboratory. Male and female mice (8 – 20 weeks old, age- and sex-matched between experimental conditions) were used in all experiments. All mice were maintained in the animal facilities at the Johns Hopkins University under specific-pathogen-free conditions and following institutional animal use and care guidelines. Mice were exposed to a 14 h/10 h light/dark cycle and fed ad libitum. The room temperature and humidity were maintained and monitored. The animal protocols were approved by the Johns Hopkins University Animal Care and Use Committee (protocols MO22M15, MO19M71).

### Primary T cell cultures

CD8^+^ T cells were isolated from the spleens and lymph nodes of 8- to 20-week old age- and sex-matched C57BL/6 mice using the MojoSort mouse CD8^+^ T cell isolation kit (480035), according to the manufacturer’s instructions. CD8^+^ T cells were activated using plate-bound anti-CD3 (5 μg/ml; InVivoMab anti-mouse CD3, BioXCell, BE0002) and soluble anti-CD28 (0.5 μg/ml; InVivoMab anti-mouse CD28, BioXCell, BE0015) in T cell culture medium (TCM): RPMI 1640 media (Invitrogen) supplemented with 10% fetal calf serum (Gibco), 4 mM L-glutamine, 100 U/ml penicillin-streptomycin, 100 U/ml IL-2 (Peprotech), 55 μM beta-mercaptoethanol in a humidified incubator at 37°C, atmospheric oxygen supplemented with 5% CO_2_. Cells were stimulated for 48 h, then expanded in IL-2 only. For the generation of T_m_ cells, IL-2 was replaced by 100 U/ml IL-15 after 72 h, and cells were maintained in IL-15 only for a further 3 days. For OT-I cultures, single-cell suspensions of splenocytes were stimulated for 48 h in IL-2-containing media as above in the presence of SIINFEKL peptide (100 ng/ml). 48 h after activation, media was replaced with TCM containing IL-2 only.

### Cell lines

The mouse E.G7 lymphoblast cell line expressing OVA (EL4-OVA) was purchased from the American Type Culture Collection (ATCC no. CRL-2113; RRID: CVCL_3505). B16-OVA melanoma cells were a kind gift from Prof. Jonathan Powell. Cells were maintained in RPMI 1640 medium (EL4-OVA) or DMEM (B16-OVA) supplemented with 10% fetal calf serum, 2 mM glutamine, 100 U/ml penicillin-streptomycin, and 55 μM beta-mercaptoethanol (EL4-OVA) or 1 mM sodium pyruvate (B16-OVA), in a humidified incubator at 37°C, atmospheric oxygen supplemented with 5% CO_2_. Platinum-E (Plat-E) retroviral packaging cells were cultured in DMEM (ThermoFisher Scientific) with 10% FCS.

### Drug and metabolite treatments

All drugs and supplemented metabolites were added at initiation of cysteine starvation, i.e. 24 h before final analyses, unless otherwise stated. L-buthionine sulfoximine (0.5 mM), GSH (1 mM) were from Sigma, elesclomol (10 – 100 nM) was from MedChemExpress, aminooxyacetic acid hydrochloride (AOAA; 10 – 50 μM) was from Cayman Chemical, and GGsTop (1 – 100 μM) was from Tocris.

### T cell exhaustion by repetitive stimulation model

Isolated CD8^+^ T cells were activated as above, with anti-CD3/28 + IL-2 for 48 h, and cultured in IL-2 only for a further 48 h. On day 4 of culture, cells were split into Acute and Chronic groups. Acute cells were re-cultured with IL-2 only every 48 h until days 6 – 10. Chronic cells were restimulated with anti-CD3/28 + IL-2 every 48 h until days 6 – 10^40,41^.

### *Listeria monocytogenes* infection studies

CD45.1^+^ recipient mice were injected i.v. with a sublethal dose of 5x10^6^ colony-forming units (CFUs) of recombinant *L. monocytogenes*-expressing OVA (*Lm*OVA). One day after infection, 1x10^5^ OT-I CD45.2^+^ control, NFS1-deficient, or CTH-deficient, CD8^+^ T cells, isolated from congenic CD45.2^+^ donor mice, were transferred i.v. into the infected mice. Peripheral blood was sampled daily from day 3 – day 8 of infection, and subjected to flow cytometry analysis.

### Mouse melanoma model

CD45.1^+^ recipient mice were shaved and 1x10^6^ B16-OVA cells in 100 μl PBS were injected into the right flank. Six - seven days after tumor injection, when tumors were palpable, 5x10^5^ OT-I CD45.2^+^ CD8^+^ control, NFS1-deficient, GCLC-deficient, or NFS1-overexpressing T cells, isolated from congenic CD45.2^+^ donor mice, were transferred i.v. into the tumor-bearing mice. Tumor sizes were measured with callipers at least every second day. The maximal allowed tumor size was 20 mm in diameter and animals were humanely euthanized if this size was reached, or if there was tumor ulceration or bleeding. Mouse body condition was monitored over the whole experimental period. At the indicated time point, mice were humanely euthanized and the tumors were excised. Tumors were digested by shaking for 1 h at 37°C with 1 mg/ml collagenase IA (Sigma, C9891) and 50 μg/ml DNase I (Roche, 10104159001). Digested tumors were passed through a 70 μm and then a 100 μm filter, and the lymphocyte fraction was obtained using a Percoll gradient.

### MC38 colon cancer model

1x10^6^ MC38 cells in 100 μl PBS were subcutaneously injected into the right flanks of BALB/c mice. Every second day from day 7, mice were treated i.p. with ICB (α-PD-1 + α-CTLA4), until harvest on day 19.

### CRISPR/Cas9 deletion

Primary mouse CD8^+^ T cells were electroporated with gRNA-Cas9 ribonucleoproteins (RNPs) using the Amaxa 4D-Nucleofector system (Lonza), 48 h after initial activation. RNPs were prepared by complexing 180 pmol of gRNAs (IDT) with 60 pmol of Cas9 nuclease V3 (IDT). Electroporation was performed in P4 Primary Cell solution using the program CM137. For CD45.2^+^ CD8^+^ OT-I cells adoptively transferred into *Lm*OVA-infected mice, cells were electroporated on day 0 using the program DS137. The nucleotide sequences for gRNAs are: control/non-targeting, GCGAGGTATTCGGCTCCGCG; NFS1, ACCTCTCACATGGACGTAC; GCLC, TACATGATCGAAGGAACGCC; ISD11 (*<u>Lyrm4</u>*), CACAAGTTTTAGATCTGTAC; CTH, AAGATGGGTAATCGTAATGG; GLRX1, TAAATAATCTTGAATCGCAC; GLRX2, GAGTTGGATATGCTGGAATA; CTACAACGTGCTGGACGACC. GLRX3, TTCATTAGTACTAGGAGGAA; GLRX5,

### Western blot analysis

For western blot analysis, cells were lysed in 1X cell lysis buffer (Cell Signaling Technology), containing 1 mM phenylmethyl sulfonyl fluoride (PMSF). Samples were frozen and thawed, then centrifuged at 18,000 g for 10 min at 4°C. Cleared protein lysate was denatured with LDS loading buffer (1X loading dye, 1 mM DTT) for 15 min at 35°C and loaded on precast 4 – 12% Bis-Tris protein gels (Life Technologies). Proteins were transferred onto nitrocellulose membranes using the iBLOT2 system (Life Technologies), according to the manufacturer’s protocols. Membranes were blocked for 1 h at room temperature with 5% w/v blotting buffer (Biorad) in Tris-buffered saline with 0.1% v/v Tween-20 (TBS-T), then incubated with primary antibodies in 5% w/v blotting buffer in TBS-T overnight at 4°C, with shaking. All primary antibody incubations were followed by at least three washes with TBS-T, then incubation with secondary horseradish peroxidase-conjugated antibody (goat anti-rabbit IgG, 31460 or goat anti-mouse IgG, 31430, 1:10,000; Pierce/Thermo Scientific) in 5% w/v blotting buffer, for at least 1 h at room temperature. After a further three washes with TBS-T, membranes were incubated for 10 min with SuperSignal West Pico or Femto chemiluminescent substrate (Pierce), and visualized using a ChemiDoc imaging system. Antibodies used were: GCLC, CTH, CTU2, ABCE1, aconitase-2, LYRM4, total OXPHOS rodent WB antibody cocktail (abcam), β-actin, p21, TOM20, GLRX1, GLRX3 (Cell Signaling Technology), ISCU (Proteintech), NFS1 (Santa Cruz), CHAC2 (Genetex), GLRX5, and GLRX2 (Novus). All primary antibodies were used at a dilution of 1:1000, except CTH and LYRM4 (1:500).

### Flow cytometry

Extracellular staining was performed in 2% FBS/PBS for 30 min, and dead cells were excluded with a Live/Dead fixable dead cell stain kit (Thermo Fisher Scientific). For samples from in vivo models, i.e. *Lm*OVA infection and B16-OVA melanoma models, F_c_ block (1:200) was added at the same time as the live/dead stain. For intracellular cytokine staining, cells were restimulated with PMA (50 ng/ml; Sigma) and ionomycin (500 ng/ml; Sigma) in the presence of brefeldin A (0.1%; Biolegend) for 5 h before extracellular staining, followed by fixation with Cytofix Cytoperm (BD Biosciences). For intracellular staining, antibodies were diluted 1:800 in permeabilization buffer and incubated with the cells overnight at 4°C. For surface staining, the following antibodies were used (1:400 unless stated otherwise): anti-CD8a (clone 53-6,7, Biolegend 100738, 100743, 100706), anti-CD44 (clone IM7, Biolegend, 103040, 103006, 103008, 103012), anti-CD62L (clone MEL-14, Biolegend, 104424, 104438, 104453), anti-CD69 (clone H1.2F3, Biolegend, 104508), anti-CD45.1 (clone A20, Biolegend, 110739, 110718), anti-CD45.2 (clone 104, Biolegend, 109832, 109808, 109806), anti-CD39 (clone Duha59, Biolegend, 143812), anti-TIM-3 (clone B8.2C12, Biolegend, 1340100), anti-PD-1 (clone 29F.1A12, Biolegend, 135225), anti-PD-L1 (clone 10F.9G2, Biolegend, 124312). For intracellular staining, the antibodies were: anti-IFNγ (clone XMG1.2, Biolegend, 505813, 505810), anti-TNF (clone MP6-XT22, Biolegend, 506318, 506304, 506306), anti-IL-2 (clone JES6-5H4, Biolegend, 503808), anti-GzmB (clone QA16A02, Biolegend, 372206), anti-NFS1 (clone B-7, Santa Cruz, sc-365308), anti-NRF2 (clone D1Z9C, CST, 14409). EdU incorporation was measured using the Click-iT EdU Alexa Fluor 647 flow cytometry assay kit according to the manufacturer’s instructions (Thermo Fisher Scientific). DNA content was measured using FxCycle Violet (1:20,000) in the final cell suspension, without washing. Cell proliferation was quantified using the CellTrace Violet (CTV) Cell Proliferation kit for flow cytometry (ThermoFisher Scientific, 34557); briefly, isolated CD8^+^ T cells were pre-stained with CTV (1:1000) for 30 min at 37°C, then washed, and activated as described with anti-CD3/28 + IL-2. Flow cytometry analyses were performed on the FACSymphony (BD Biosciences) or Cytek Northern Lights, and then analyzed with FlowJo 10 Software (TreeStar). CellROX, MitoTracker Green (50 nM), labile iron (Fe^2+^; 5 μM), C11-BODIPY, and BODIPY stainings were performed in live cells, for 30 min at room temperature in PBS, and analyzed immediately.

### Cytolysis assay

OT-I CD8^+^ T cells were stimulated with plate-bound anti-CD3 (5 μg/ml; InVivoMab anti-mouse CD3, BioXCell, BE0002) and soluble anti-CD28 (0.5 μg/ml; InVivoMab anti-mouse CD28, BioXCell, BE0015) in RPMI 1640 media (Invitrogen) supplemented with 10% fetal calf serum (Gibco), 4 mM L-glutamine, 100 U/ml penicillin-streptomycin, 100 U/ml IL-2 (Peprotech), 55 μM β-mercaptoethanol in a humidified incubator at 37°C, atmospheric oxygen supplemented with 5% CO_2_. Cells were stimulated for 48 h, then expanded in IL-2 only for a further 72 h. Cells were then cultured in full or cysteine-free medium overnight, then washed and plated with CellTrace Violet pre-stained EL4-OVA cells over a dilution series, in full or cysteine-free medium. EL4-OVA cells, without T cells, were also plated in full or cysteine-free medium. The percentage of dead EL4-OVA cells was assessed after 6 h, by flow cytometry.

### Seahorse analysis

One day prior to the assay, 200 μl of XF calibrant was added to each well of a 96-well Seahorse calibrant plate. 2x10^5^ cells per well were plated on a Seahorse 96-well cell culture plate in 40 μl of the appropriate medium (full medium, or without the indicated amino acids), and the plate was spun at 300 g for 3 min. Then, 100 μl of the appropriate medium was added to the wells for a final volume of 140 μl. OCR was measure using the Seahorse analyzer maintained at 37°C. The complex V (ATP synthase) inhibitor oligomycin (10 μM), mitochondrial membrane ionophore FCCP (5 μM), complex I inhibitor rotenone (100 nM), and complex III inhibitor antimycin A (1 μM) were injected as indicated.

### ATP assay

1x10^5^ cells were lysed by boiling in H_2_O, and analyzed using the ATP determination kit (Thermo Fisher Scientific), according to the manufacturer’s instructions.

### Glutathione quantification

GSH and total glutathione (GSH + GSSG) were quantified using the GSH-Glo glutathione assay (Promega), according to the manufacturer’s instructions. 2x10^5^ cells per condition were assayed in a 96-well U-bottomed plate. 50 μl of 1X GSH-Glo reagent was added to each well, and the plate was incubated at room temperature for 30 min. To detect total glutathione, the reducing agent TCEP was added to 1X GSH-Glo reagent at a concentration of 1 mM, to reduce GSSG to GSH. 50 μl of reconstituted luciferin detection reagent was added to each well, and the plate was incubated in the dark for 15 min. 90 μl of each sample was transferred to a white, opaque luminometer plate and luminescence was measured using a plate reader (Details). GSH and total glutathione concentrations were calculated from a GSH standard curve, and GSSG levels were calculated by subtracting GSH from total glutathione.

### GGT Assay

1x10^6^ cells were lysed in GGT Assay Buffer, and analyzed using the GGT colorimetric assay kit (Abcam), according to the manufacturer’s instructions.

### Metabolite extraction

1x10^6^ cells were centrifuged for 5 min at 500 g, 4°C. 5 μl of the supernatant was removed into a new 1.5 ml Eppendorf tube, and 95 μl of 80% ice-cold extraction buffer (80%) methanol was added. The pellet was washed with ice-cold PBS and centrifuged for a further 5 min 500 g, 4°C. Samples were extracted in 100 μl 80% methanol. Cell and supernatant samples were immediately frozen at -80°C. For NMR analyses, 40x^6^ cells were extracted in 1 ml extraction buffer. For metabolite extraction from ISF, tumor sections were placed on a 1 nm mesh in a 5 ml tube, and centrifuged for 5 min, 500g, at 4°C. 2 μl of the flow through was collected, avoiding any pelleted cells. 50 μl methanol (80%) was added, and samples were frozen at -80°C. Before measurement, samples were centrifuged for 10 min at maximum speed, 4°C, and transferred into new 1.5 ml Eppendorf tubes. 40 μl of sample was added to a plastic LC-MS vial, and subjected to LC-MS analysis.

### LC-MS metabolomics analysis and stable isotope tracing

Chromatographic separation was performed on an Agilent 1290 Infinity II UHPLC system using a Waters Atlantis Premier BEH Z-HILIC VanGuard FIT column (1.7 μm, 2.1 mm x 100 mm). Mobile phase A was 20 mM NH_4_CO_3_, 5 μM medronic acid in water. Mobile phase B was 10% mobile phase A + 90% acetonitrile. The gradient profile was: 0 min, 95% B, 0.18 ml/min; 14 min, 55% B, 0.18 ml/min; 15 min, 20% B, 0.18 ml/min; 19 min, 20% B, 0.18 ml/min; 19.50 min, 95% B, 0.18 ml/min; 22 min, 95% B, 0.18 ml/min; 25 min, 95% B, 0.4 ml/min; stop time, 29 min. Column temperature was 40°C and autosampler temperature was 4°C. Injection volume was 3 μl. The LC system was coupled to a Bruker timsTOF-Pro mass spectrometer (MS) equipped with an ionBooster electrospray ionization (ESI) source. The mass spectrometer was operated in both positive and negative modes with tims on. The mass axis was calibrated at the beginning of every sample run. Data were acquired with the Compass Hystar software. Metabolite peak areas were identified and determined using Metaboscape and TASQ software (versions 2022 – 2024b; Bruker).

### NMR analysis

NMR spectra were acquired using Bruker Avance II or Bruker NEO 600 MHz (^1^H) spectrometers, equipped with 5 mm triple-resonance cryogenic probes and *z*-axis pulsed-field gradients. In-house variations of ^1^H-^13^C HSQC pulse sequences with aliphatic ^13^C selectivity, sensitivity-enhanced quadrature detection and gradient-based coherence selection were employed. Constant-time and real-time versions were used for the ^13^C-Cys samples and Cys, GSH controls, respectively. Typical experimental parameters were: (1) General: temperature 25°C, number of scans 16 or 32/FID, recycle delay between scans 1.2 s, constant-time delay period 28 ms, total time 2.5 – 5 h per experiment, (2) ^1^H: center of spectrum 4.77 ppm (H_2_O frequency), spectral width 20 ppm, acquisition time (t_2_max) 10 ms, 2750 data points/FID, (3) ^13^C: center of spectrum 50.5 ppm, spectral width 80 ppm, acquisition time (t_1_max) 16 ms, 390 data points, and a 350 μs Reburp pulse was used to select for aliphatic ^13^C nuclei spanning 0 to 90 ppm. 2D data were processed using nmrPipe software and analyzed using Sparky. 1D spectra were processed and analyzed via Topspin 4.3.0 software.

### Retroviral overexpression of NFS1

Murine stem cell virus (MSCV)-based plasmids were used to overexpress CD90.1/Thy1.1 (as a control plasmid with a marker for detection by flow cytometry; MSCV-Bbsl-2A-Thy1.1) or NFS1 and CD90.1 together (MSCV-NFS1-2A-Thy1.1). In the NFS1 plasmid, the Thy1.1 sequence was added immediately after the NFS1 cDNA sequence followed by 2A self-cleavage peptide, both to serve as a marker for detection by flow cytometry, and to indicate successful expression of NFS1. Plasmids contained an ampicillin resistance gene. Plasmids were transformed into Stable Competent *E. coli* cells (New England Biolabs) via heat shock. Successfully transformed bacteria were selected by growth in LB broth with ampicillin, and plasmid DNA was extracted using a Qiagen MiniPrep kit. Plat-E cells were cultured at a density of 0.75x10^^6^ cells/ml in 2 ml of Opti-MEM reduced serum medium with 10% FCS overnight before transfection with the plasmid of interest. On the day of transfection, the transfection mixture was prepared by making tubes A and B as follows: tube A was prepared with 210 μl Opti-MEM (no FCS), 6.19 μl p3000, 0.9 μg transfer plasmid (MSCV-Bbsl-2A-Thy1.1 or MSCV-NFS1v2-2A-Thy1.1), and 0.9 μg pCL-Eco helper plasmid, and tube B with 210 μl Opti-MEM (no FCS) and 7.04 μl Lipofectamine 3000 (all quantities per reaction). Tubes A and B were vortexed, combined, and incubated at room temperature for 15 min. 850 μl of media was removed from each well of Plat-E cells, and after 15 min, the transfection mixture was added to the appropriate wells. Plat-E cells were incubated at 37°C for 6 hours, at which point the media was removed and replaced with 2 ml Opti-MEM (10% FCS). The culture media was collected 48 h later, fresh media was added to the cells, and this media was collected after a further 24 h and combined with the previously collected media. This was filtered using a 0.45 μm filter and 1/3 volume retrovirus-X concentrator was added. This was mixed by inversion, incubated at 4°C overnight, then centrifuged at 1500 g, 4°C, for 45 min. The supernatant was removed and the pellet of virus was resuspended in TCM at 1/10 – 1/100 of the original volume, before freezing at -80°C (for storage) or addition to T cells. Retroviral transduction was performed using primary CD8^+^ T cells that had been activated for 20 – 24 h with αCD3/28 + IL-2. T cells were plated at a density of 2 – 3x10^6^/ml in TCM. Polybrene (4 μg/ml) and the concentrated virus were added to the appropriate wells, and the cells were spinoculated at 2000 g, 32°C, for 90 min. The plate was then incubated at 37°C for 6 h. T cells were then replated on an αCD3-coated plate, with αCD-28 + IL-2. After 24 h, T cells were replated with IL-2, and cultured as described above.

### scRNAseq analysis

Single cell RNA-sequencing data from hepatocellular carcinoma patients were retrieved from the European Genome-Phenome Archive (EGAC00001000551) containing immune cells isolated from peripheral blood and tumor or liver biopsies^43^. Deposited BAM files aligned to GRCh38 were downloaded and read counts for individual cells obtained using ‘featureCounts’^79^ from Rsubread v.2.18.0. The resulting raw read count matrix of barcodes corresponding to cells and features corresponding to detected genes were processed, analyzed and visualized in R v. 4.3.1^80^ using Seurat v.5^81^ with default parameters in all functions, unless specified. Filtered samples were normalized and scaled using ‘NormalizeData’ and ‘ScaleData’ to identify highly variable genes using ‘FindVariableFeatures’, blacklisting genes associated with tissue processing based on annotation available in SignatuR v. 0.2.1^82^. Concomitantly, samples were regularized using negative binomial regression (SCTransform)^83^, batch corrected with Harmonyv. 1.2.0^84^ using ‘RunHarmony’ with 30 principal components calculated on the SCT assay using ‘RunPCA’. Batch corrected dimensionality reductions were used to identify neighbours using ‘FindNeighbors’, cluster cells using ‘FindClusters’ and calculate a Uniform manifold approximation and projection (UMAP)^85^ using ‘RunUMAP’. Resolution for cell clustering was determined by evaluating hierarchical clustering trees and silhouette scores at a range of resolutions (0 - 1.2, with 0.05 step increments) with Clustree v. 0.5.1^86^ and the ‘silhouette’ function in cluster v. 2.1.6^87^, selecting a value inducing minimal cluster instability and a high proportion of silhouette widths above 0. Marker genes associated with clusters were identified by using a Wilcoxon Rank Sum test with ‘FindMarkers’ from Seurat v.5^83^. Cell type scores were calculated using scGate v.1.6.2^88^.

### Statistical analysis

Statistical analysis was performed using Prism 10 software (GraphPad). Results are represented as mean ± SEM, unless otherwise indicated. *n* represents individual biological replicates. Comparisons of two groups were calculated using an unpaired two-tailed Student’s *t*-test. The selection of sample size was based on extensive experience with metabolic, immunologic, and tumor assays, in vitro and in vivo.

---

**Supplementary Figure 1.**
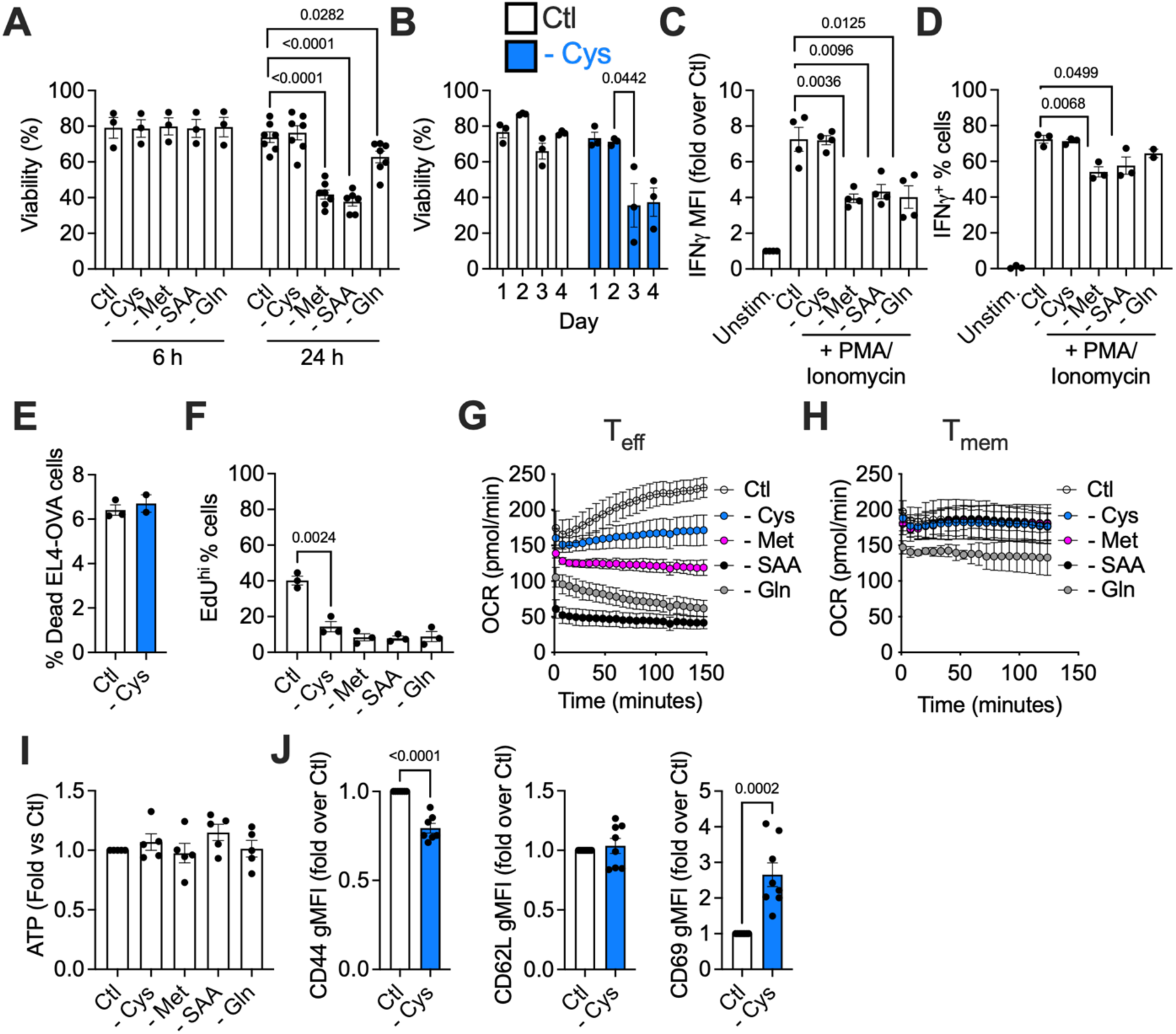
Enhanced cytokine production is specific to cysteine starvation. (A – D, F – J) Isolated, activated (with αCD3/28) CD8^+^ T cells were starved of cysteine (- Cys), methionine (- Met), both cysteine and methionine together (- SAA), or glutamine (- Gln) for 6 - 24 h (from day 5 to day 6 of culture). In (B), cells were starved of cysteine for the days indicated. (A, B) Viability, (C) IFNγ levels, (D) frequency of IFNγ^+^ cells, in fed and starved cells. In (C, D) cells were restimulated for 6 h with PMA/ionomycin, concurrent with starvation. (E) EL4-OVA lymphoma cells were fed or starved of cysteine for 6 h, and viability was measured by flow cytometry. (F) Percentage of EdU^hi^ cells, in fed and starved cells, starved for 6 h to maintain viability in all conditions. (G, H) OCR by Seahorse analysis in T_eff_ (G) and T_mem_ (H) cells, from initiation of starvation. (I) ATP levels by luminescence assay, in cells starved for 6 h. (J) Surface staining of CD44 (left), CD62L (middle), and CD69 (right) of cysteine-starved cells (24 h).

**Supplementary Figure 2.**
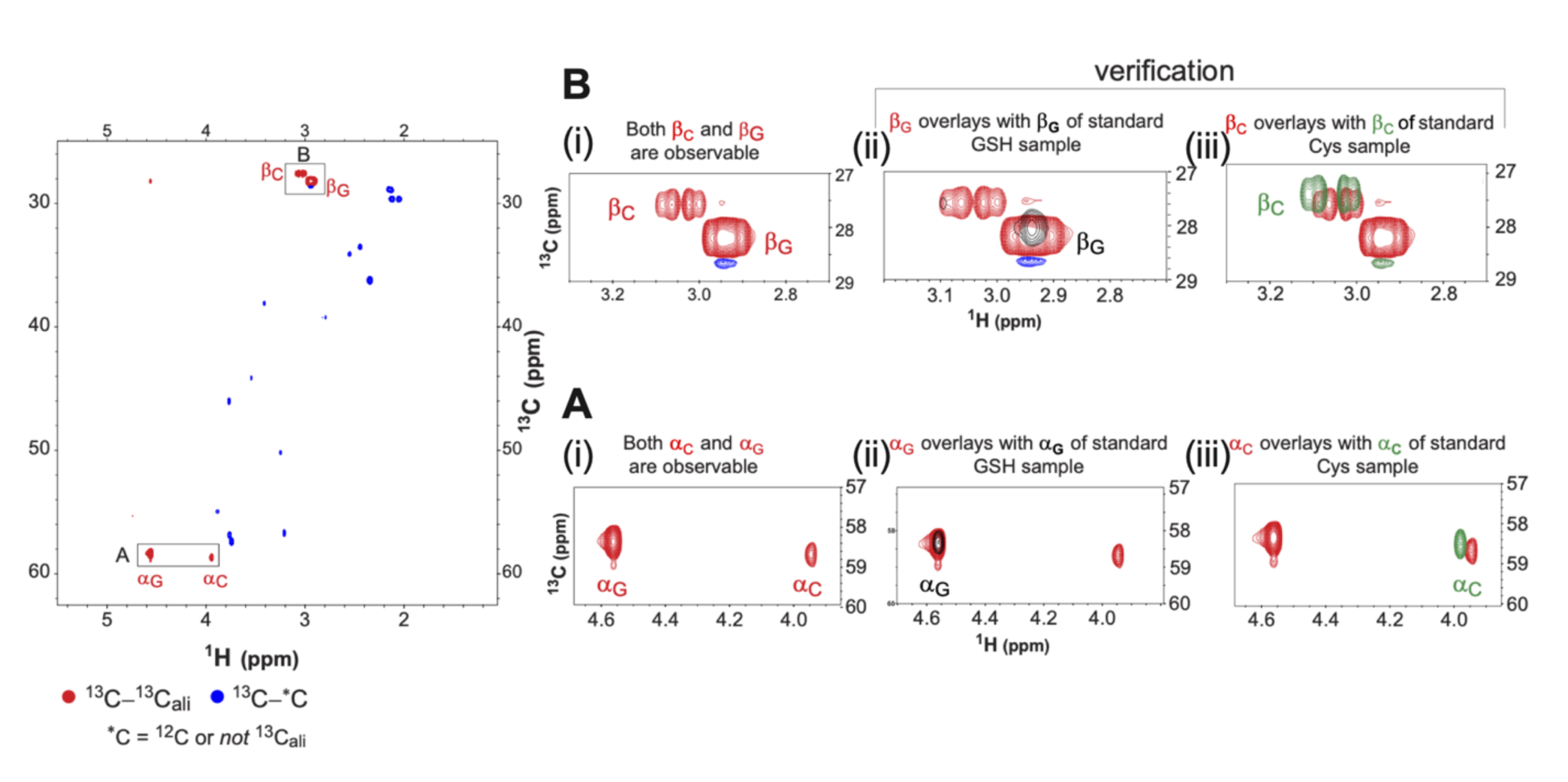
Verification of NMR peaks. Two-dimensional (2D) NMR ^1^H-^13^C heteronuclear single quantum coherence (HSQC) experiment which correlates a ^1^H nucleus with its attached ^13^C nucleus (^1^H:^13^C). The NMR chemical shifts of ^1^H (ppm) and ^13^C (ppm) are represented along the *x* and *y* axes of the 2D spectrum respectively, and the ^1^H:^13^C correlation is manifested as a peak at the intersection of the chemical shift coordinates. The spectrum was acquired in constant-time (CT) mode such that ^1^H:^13^C correlations from a ^1^H-^13^C-^13^C_ali_ (ali = aliphatic) linkage has the opposite phase (in red) relative to ^1^H-^13^C-*C linkages (in green), where *C is either ^12^C or a non-aliphatic carbon (e.g. aromatic or CO or CO_2_). As cysteine is of the form Hα-Cα-C-(H1, H2), uniformly ^13^C-labeled Cys in the sample gives rise to one Hα:Cα peak (A(i), alpha carbons, α_C_), and two Hβ_1,2_:Cβ peaks (B(i), beta carbons, β_C_). The entire ^13^C-Cys carbon skeleton is incorporated into GSH since the corresponding Hα:Cα (α_G_) and Hβ_1,2_:Cβ peaks in GSH (β_G_) have the same phase as ^1^H-^13^C-^13^C_ali_ linkages. For GSH, Hβ1 and Hβ2 chemical shifts are very similar, resulting in near coalescence of the Hβ1:Cβ and Hβ2:Cβ peaks. Blue peaks arise from ^1^H-^13^C correlations from other species in solution, at the 1% natural abundance of ^13^C, and as a result are almost exclusively ^1^H-^13^C-^12^C linkages, and therefore of opposite phase to the Cys and GSH peaks. Control ^1^H-^13^C HSQC spectra of Cys and GSH standards were separately acquired to confirm that the peaks of interest were from Cys and GSH (verification, A(ii, iii), B(ii, iii)). Overlays of the ^13^C-Cys-incubated sample spectrum (red) with the control GSH spectrum (black) are shown in A(ii) and B(ii), and with the control Cys spectrum (green) in A(iii) and B(iii). The control samples were not ^13^C-labeled, and so the spectra relied on the 1% natural abundance of ^13^C, i.e. of the form ^1^H-^13^C-^12^C, resulting in a slight isotope shift between ^1^H:^13^C-^13^C and ^1^H:^13^C-^12^C peaks, and likely causing complete collapse of the two H1,2 protons in GSH. The overlaid spectra clearly establish the identities of the representative Cys and GSH peaks in the ^13^C-Cys-incubated sample spectrum.

**Supplementary Figure 3.**
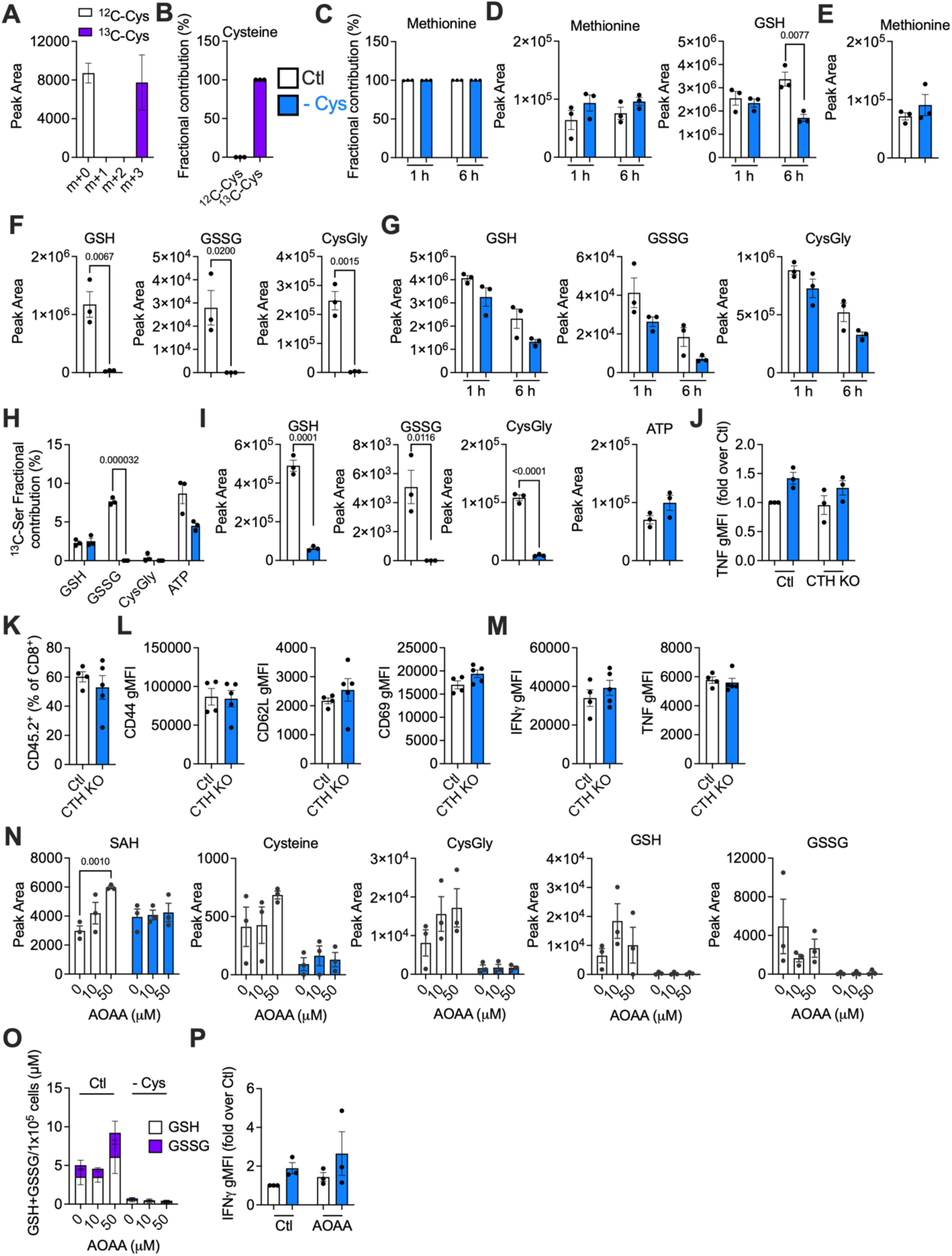
Transsulfuration does not replenish cysteine or GSH. (A) Pool sizes of ^12^C-Cys (unlabelled, m+0) and ^13^C-Cys (labelled, m+3) in CD8^+^ T cells incubated with ^12^C- or ^13^C-Cys for 24 h. (B) Fractional contribution of ^12^C- or ^13^C-Cys to cysteine after 24 h. (C – M) Cells were fed (Ctl) or starved of cysteine (- Cys) for 1 – 24 h as indicated. (C) Fractional contribution of ^13^C-Met to methionine after 1 - 6 h. (D) Pool sizes of methionine and GSH in cells incubated with ^13^C-Met for 1 - 6 h. (E, F) Pool sizes of methionine (E), GSH, GSSG, and CysGly (F) in cells incubated with ^13^C-Met for 24 h. (G) Pool sizes of GSH, GSSG, and CysGly in cells incubated with ^13^C-Ser for 1 - 6 h. (H) Fractional contribution of ^13^C-Ser to the indicated metabolites after 24 h. (I) Pool sizes of GSH, GSSG, CysGly, and ATP in cells incubated with ^13^C-Ser for 24 h. (J) TNF after PMA/ionomycin restimulation in fed and starved (24 h) WT or CTH KO T cells. (K – M) CTH was deleted by CRISPR/Cas9 targeting in isolated naïve CD8^+^ T cells (day 0), and 0.1x10^6^ control or CTH-deficient OT-I CD45.2^+^ CD8^+^ T cells were transferred into CD45.1^+^ mice that had been infected one day previously with 5x10^6^ LmOVA. Mice were bled daily and samples were stained for flow cytometric analysis. (K) expansion of CD45.2^+^ CD8^+^ donor T cells. (L) CD44, CD62L, CD69 expression by flow cytometry. (M) IFNγ and TNF after 5 h restimulation with PMA/ionomycin. (N – P) Cells were treated with AOAA (10 – 50 μM) for 24 h, concurrent with cysteine starvation. (N) Pool sizes of SAH, cysteine, CysGly, GSH, and GSSG in control or AOAA-treated, fed and cys-starved cells (24 h). (O) Luminescence assay for GSH and GSSG. (P) IFNγ and TNF after PMA/ionomycin restimulation.

**Supplementary Figure 4.**
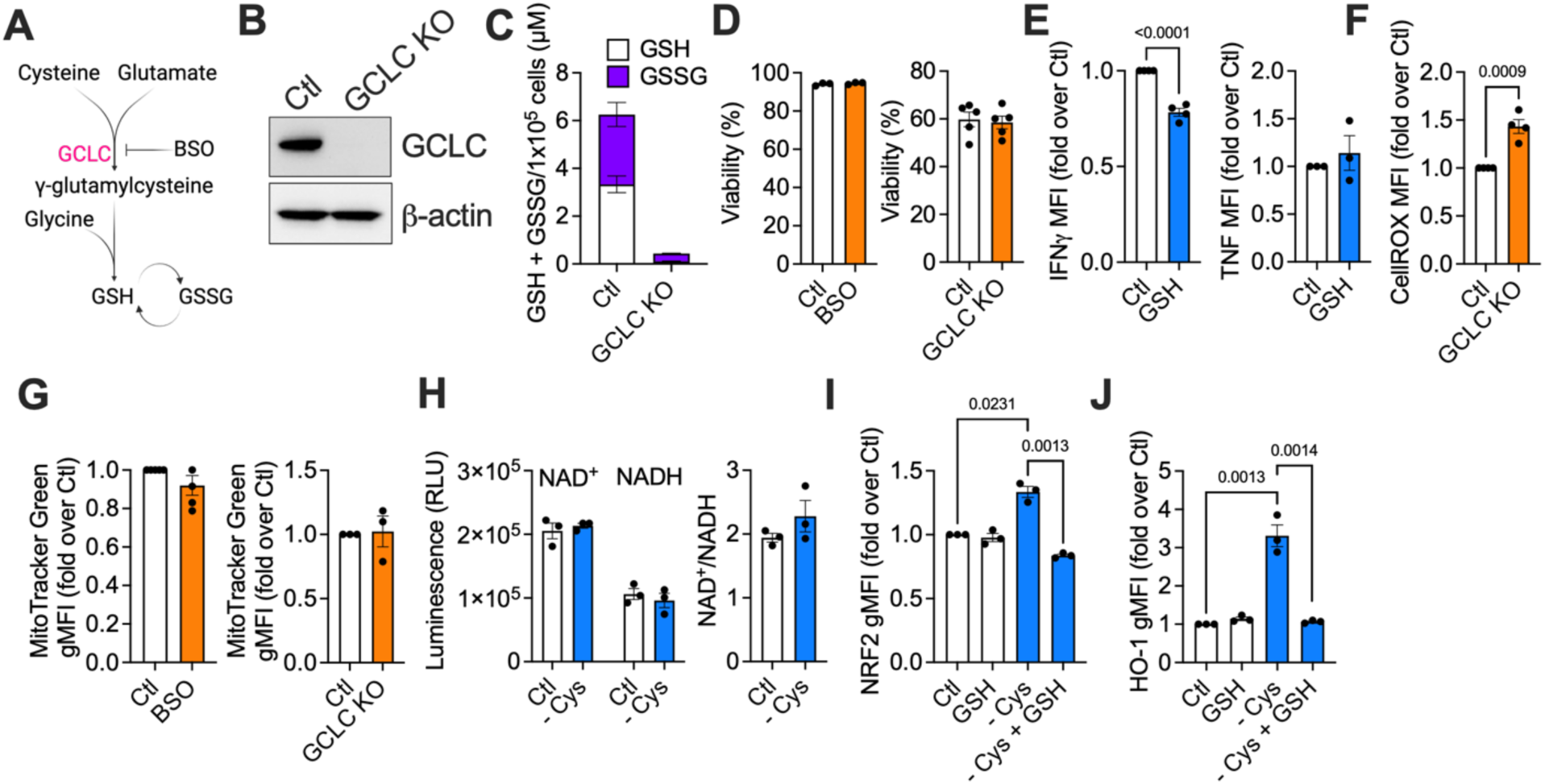
Glutathione disruption does not impact T cell viability or mitochondrial content. (A) Schematic of glutathione synthesis from cysteine, indicating action of GCLC and BSO. (B) Western blot confirming GCLC deletion. (C) GSH and GSSG by luminescence assay. (D) Viability in WT (Ctl), BSO-treated (left) or GCLC KO (right) cells. (E) Fed cells were treated with GSH (1 mM) for 24 h prior to restimulation for 5 h with PMA/ionomycin, and IFNγ (left) and TNF (right) were measured by flow cytometry. (F) CellROX staining of ROS in WT and GCLC KO cells. (G) MitoTracker Green staining for mitochondrial content in WT, BSO-treated (0.5 mM; 24 h; left), or GCLC KO (right) cells. (H) NAD^+^ and NADH levels (left), and the NAD^+^/NADH ratio (right) in fed or cys-starved (24 h) T cells. (I, J) T cells were fed or cysteine-starved (24 h), +/- GSH (1 mM, 24 h, concurrent with cysteine starvation. NRF2 (H) and the NRF2 target HO-1 (I) were measured by flow cytometry.

**Supplementary Figure 5.**
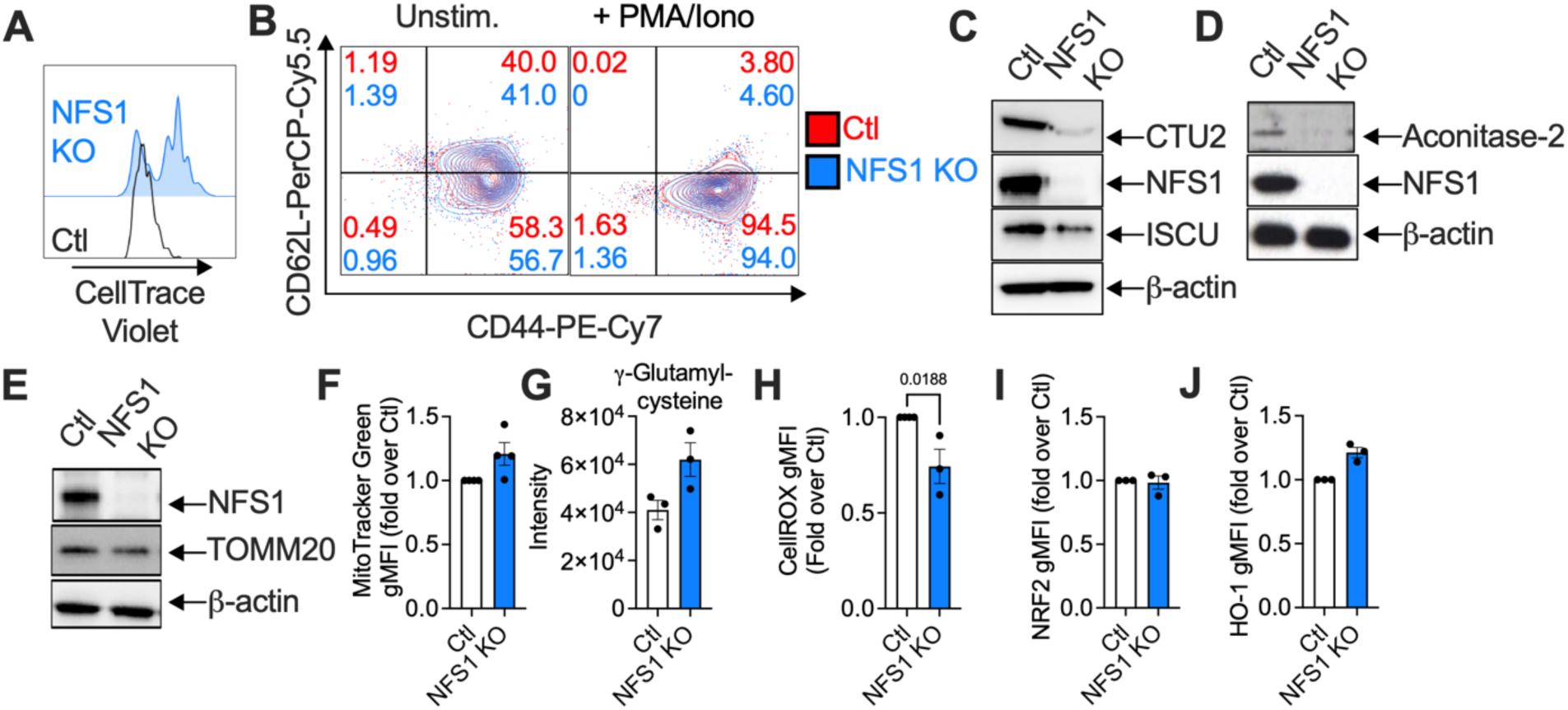
NFS1-deficient T cells have lower levels of FeS-cluster containing enzymes, but maintain mitochondrial mass. (A) NFS1 was deleted by CRISPR/Cas9 targeting in isolated naïve CD8^+^ T cells (day 0). WT (Ctl) or NFS1 KO cells were stained with CellTrace Violet, and proliferation was assessed on day 4. (B - J) NFS1 was deleted by CRISPR/Cas9 in CD8^+^ T cells after 2 days of activation. (B) Flow cytometric staining of activation markers CD44 and CD62L, in unstimulated and restimulated (with PMA/ionomycin) Ctl and NFS1 KO cells. (C – E) Western blots showing NFS1 deletion, and levels of (C) downstream NFS1 effectors ISCU (FeS cluster synthesis), CTU2 (tRNA thiolation), (D) aconitase-2 (an FeS cluster enzymes that is not part of the ETC), and (E) TOMM20, a mitochondrial protein that does not have an FeS cluster. (F) MitoTracker Green staining for mitochondrial mass, (G) γ-glutamylcysteine levels by LC-MS, (H) CellROX staining for ROS, (I) NRF2 expression, and (J) HO-1 expression, in Ctl and NFS1 KO T cells.

**Supplementary Figure 6.**
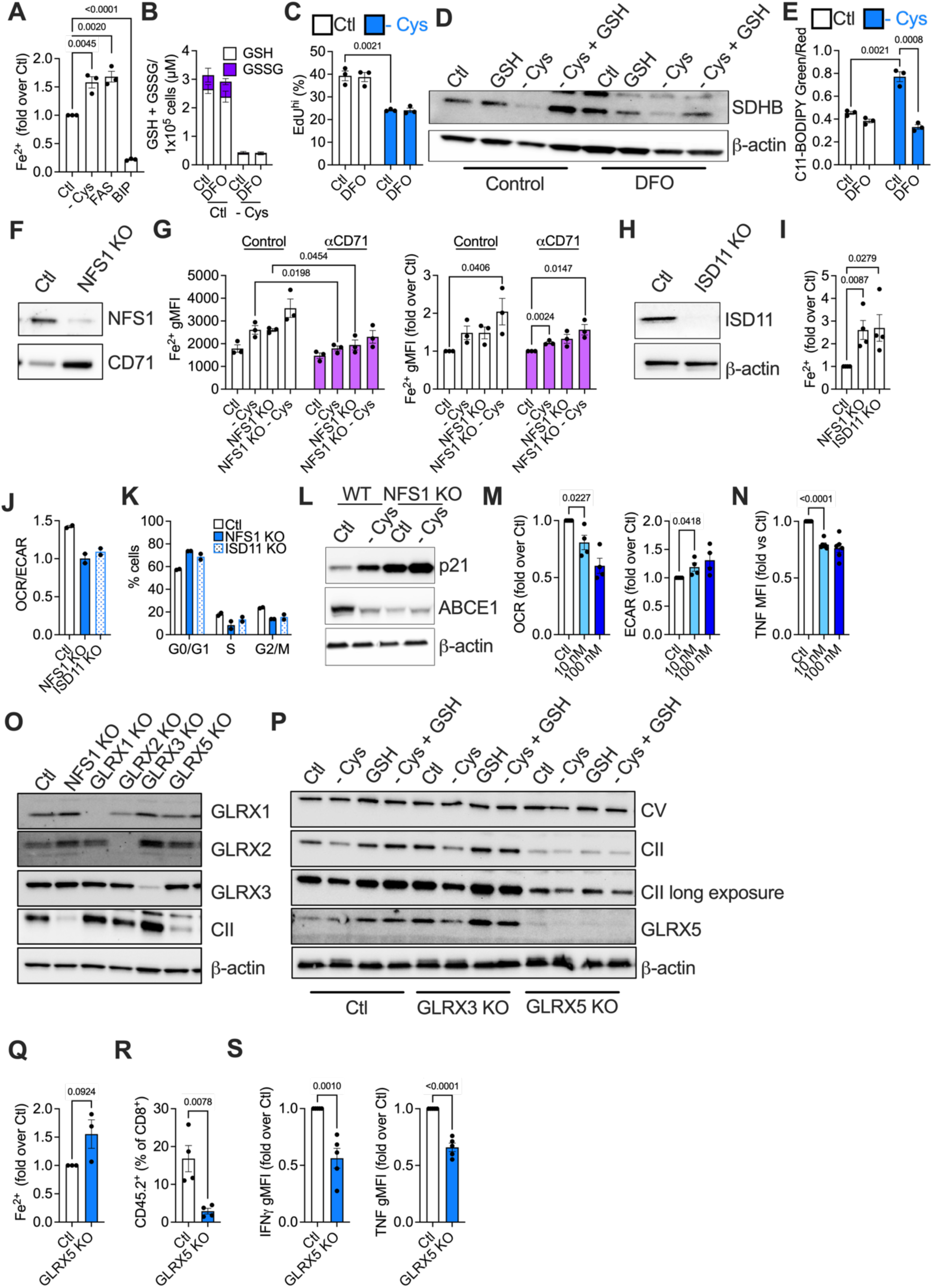
GSH stabilizes FeS clusters via GLRX5. (A) Cells were fed or cys-starved (24 h), or treated with ferrous ammonium sulfate (FAS; 100 μM) or bipyridyl (BIP; 100 μM) for 0.5 h. (B, C) Cells were fed or cys-starved for 24 h, and concurrently treated with DFO (5 μM). (B) GSH and GSSG were measured by luminescence assay. (C) Proliferation was measured by EdU staining. (D) Cells were fed or cys-starved, and treated +/- GSH (1 mM) +/- DFO (5 μM) for 24 h. SDHB (CII) was measured by western blot. (E) Cells were fed or cys-starved for 24 h, and concurrently treated with DFO (5 μM). Lipid peroxidation was measured by C11-BODIPY staining. (F - L) NFS1 or ISD11 was deleted by CRISPR/Cas9 targeting in isolated naïve CD8^+^ T cells (day 2). (F) Western blot showing NFS1 and CD71 levels. (G) Fe^2+^ was measured using BioTracker far-red labile Fe^2+^ dye. (H) Western blot showing ISD11 deletion, (I) Fe^2+^ in Ctl, NFS1 KO, and IDS11 KO cells, measured using BioTracker far-red labile Fe^2+^ dye. (J) OCR/ECAR ratio, by Seahorse analysis. (K) Cell cycle analysis by EdU/FxCycle Violet staining in NFS1 KO and ISD11 KO cells. (L) Western blot of p21 and ABCE1 in fed or cys-starved (24 h) Ctl or NFS1 KO cells (M) OCR (left) and ECAR (right) in elesclomol-treated cells. (N) TNF production in T cells treated with elesclomol (10 – 100 nM; 24 h) and restimulated for 5 h with PMA/ionomycin. (O) Western blot of GLRX1, GLRX2, GLRX3, GLRX4, GLRX5, and CII in wild type (Ctl), NFS1-, GLRX1-, GLRX2-, GLRX3-, and GLRX5-deficient cells. (P) Ctl, GLRX3 KO, and GLRX5 KO cells were fed or cys-starved, and treated +/- GSH (1 mM; 24 h). CV, CII, and GLRX5 were measured by western blot. (Q) Fe^2+^ in Ctl and GLRX5 KO cells. (R) GLRX5 was deleted by CRISPR/Cas9 targeting in isolated naïve CD8^+^ T cells, and 0.1x10^6^ control or GLRX5-deficient OT-I CD45.2^+^ CD8^+^ T cells were transferred into CD45.1^+^ mice that had been infected one day previously with 5x10^6^ LmOVA. Mice were bled daily and expansion of CD45.2^+^ CD8^+^ donor T cells was measured. (S) IFNγ and TNF in Ctl and GLRX5 KO cells (GLRX5 deleted on day 2 of culture), after 5 h restimulation with PMA/ionomycin.

**Supplementary Figure 7.**
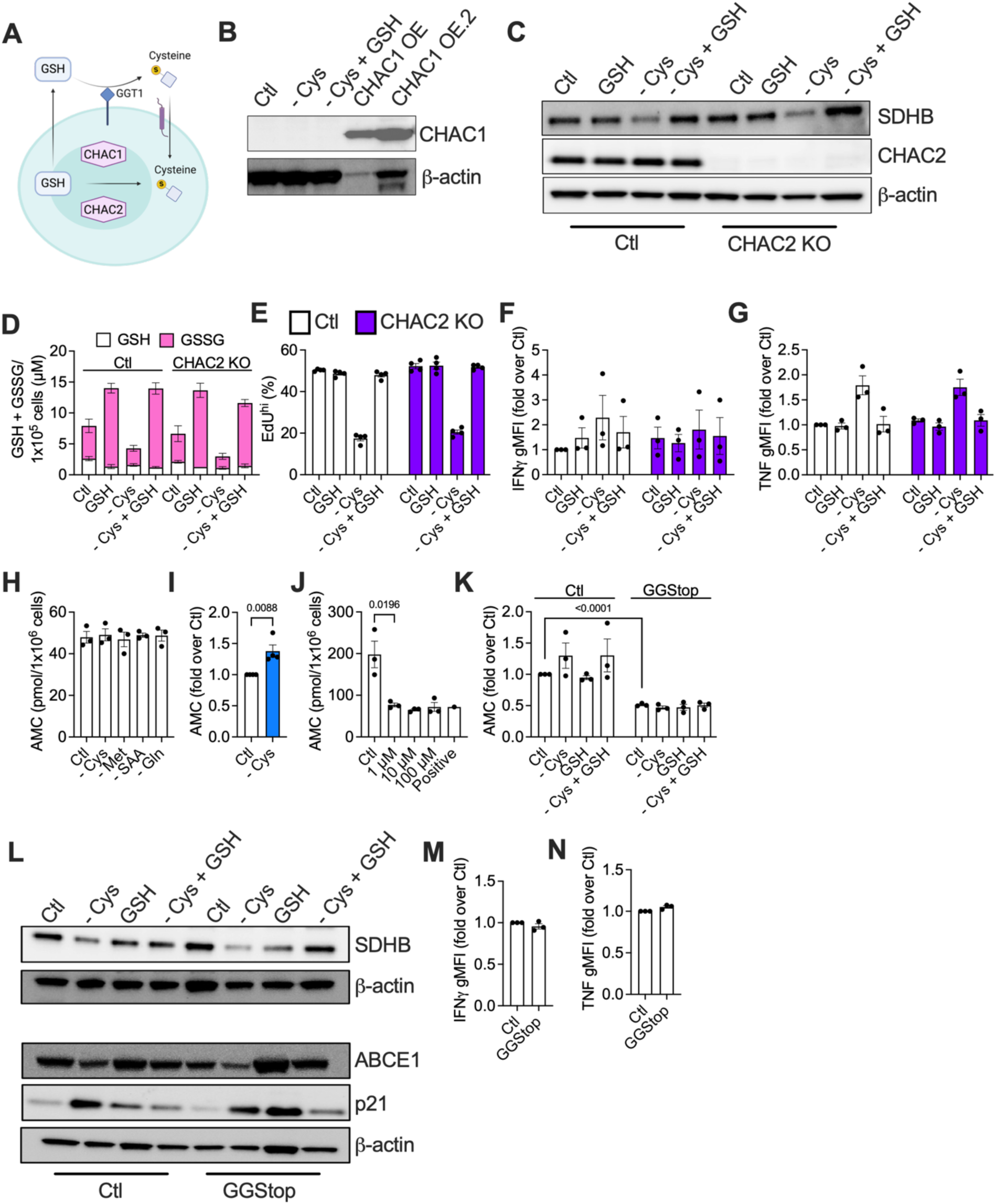
GSH is not catabolized to replenish cysteine. (A) Schematic diagram illustrating how GSH can be broken down to cysteine. (B) CHAC1 expression was measured in T cells that were fed, starved of cysteine, or starved of cysteine and supplemented with GSH (1 mM), for 24. Lysates from CHAC1 overexpressing cells (CHAC1 OE) were run alongside these samples, either at 10% of the input by protein concentration (CHAC1 OE, lane 4), or with equal protein concentration (CHAC1 OE.2, lane 5) to these test samples from fed and starved T cells (lanes 1 – 3). (C - G) Wild type (Ctl) and CHAC2 KO cells were fed or cys-starved, and treated +/- GSH (1 mM) for 24 h. (C) SDHB (CII) and CHAC2 were measured by western blot. (D) GSH and GSSG were measured by luminescence assay. (E) EdU was used to measure proliferation. (F, G) IFNγ (F) and TNF (G) after 5 h restimulation with PMA/ionomycin. (H, I) Cells were starved of the indicated amino acids for 6 h (H), or of cysteine for 24 h (I) and AMC production was measured as a readout of GGT activity. (J) Cells were treated for 24 h with the indicated doses of the GGT inhibitor GGStop, and AMC production was measured. (K) AMC production and (L) SDHB, ABCE1, and p21 levels in fed, cys-starved, and GSH-supplemented (1 mM; 24 h) cells, +/- GGStop (1 μM). (M, N) IFNγ (M) and TNF (N) after 5 h restimulation with PMA/ionomycin in cells treated with GGStop (1 μM) for 24 h.

**Supplementary Figure 8.**
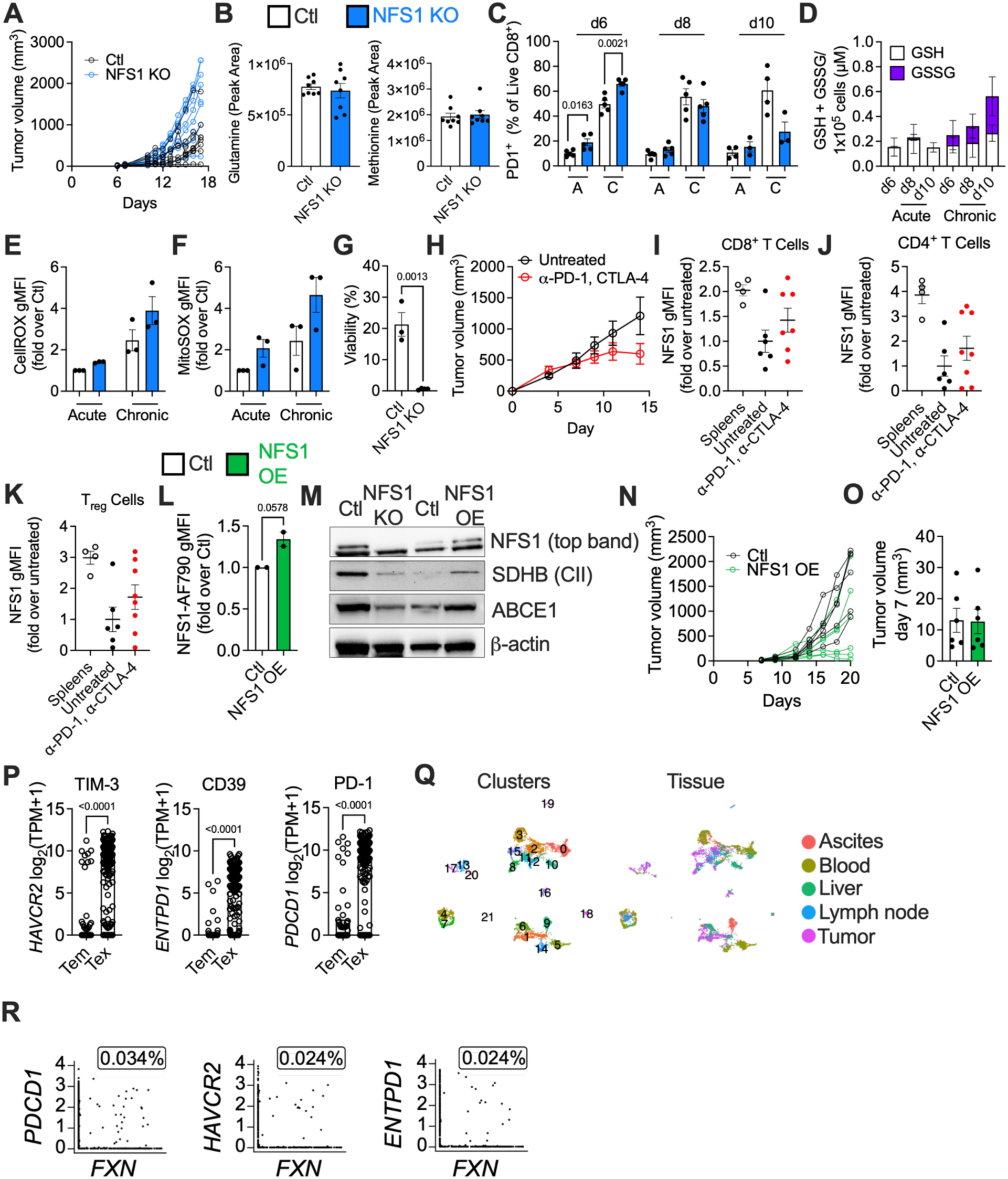
NFS1 deficiency correlates with T cell exhaustion. (A, B) 1x10^6^ B16-OVA melanoma cells were subcutaneously injected into the flanks of CD45.1^+^ recipient mice. 7 days post-implantation, when tumors were palpable, 0.5x10^6^ control (Ctl) or NFS1 KO CD45.2^+^ CD8^+^ OT-I T cells were adoptively transferred into these recipients by intravenous tail vein injection. (A), Individual tumor growth curve for each mouse. (B) Metabolites in tumor ISF, measured by LC-MS. (C – G) In an in vitro model of T cell exhaustion, control or NFS1 KO cells were initially activated for 2 days with αCD3/28 + IL-2, then cultured with IL-2 only (Acute; A) or restimulated with IL-2 + αCD3/28 (Chronic; C) every 48 h for 8 - 10 days. (C) PD1^+^ frequency in CD8^+^ T cells. (D) GSH and GSSG were measured by luminescence assay in wild type CD8^+^ T cells. (E, F) CellROX (E) and MitoSOX (F) staining in Acute and Chronic Ctl and NFS1 KO cells on day 8. (F) Viability of Chronic control or NFS1 KO cells. (H – K) 1x10^6^ MC38 colon cancer cells were subcutaneously injected into the flanks of BALB/c recipient mice. Beginning on day 7 post-implantation, mice received intraperitoneal injections of α-PD-1 + α-CTLA4 as ICB, or were untreated, every 2^nd^ day until tumor harvest on day 14. (H) Tumor growth curve of untreated mice, or mice receiving ICB. (I – K) NFS1 expression in CD8^+^ (I), CD4^+^ (J), and T_reg_ (K) subsets from spleens, untreated mice, or ICB-treated mice. (L) NFS1 expression by flow cytometry in CD8+ T cells induced to overexpress NFS1 by retroviral transduction. (M) NFS1, SDHB (CII), ABCE1, and β-actin expression by western blot in CD8+ T cells induced to overexpress NFS1 by retroviral transduction. (N, O) 1x10^6^ B16-OVA melanoma cells were subcutaneously injected into the flanks of CD45.1^+^ recipient mice. 7 days post-implantation, when tumors were palpable, 0.5x10^6^ control (Ctl) or NFS1 OE CD45.2^+^ CD8^+^ OT-I T cells were adoptively transferred into these recipients by i.v. tail vein injection. (N) Individual tumor growth curves, and (O) starting tumor volumes on day 7. (P) A previously published scRNAseq dataset of TIL subsets in human HCC patients were analyzed to determine TIM-3, CD39, and PD-1 expression in effector memory (Tem), and exhausted (Tex) TILs in these patients. (Q, R) A previously published human HCC scRNAseq analysis ws reanalyzed. (Q) 21 different cell clusters were identified (left), representing cells from ascites, blood, liver, lymph node, and tumor sites (right). (R) Co-expression of *FXN* (top) each of the exhaustion markers *PDCD1*, *HAVCR2*, and *ENTPD1* was determined, in CD4^+^ and CD8^+^ T cells.

